# Single-cell lineage and transcriptome reconstruction of metastatic cancer reveals selection of aggressive hybrid EMT states

**DOI:** 10.1101/2020.08.11.245787

**Authors:** Kamen P Simeonov, China N Byrns, Megan L Clark, Robert J Norgard, Beth Martin, Ben Z Stanger, Aaron McKenna, Jay Shendure, Christopher J Lengner

**Affiliations:** Medical Scientist Training Program, Perelman School of Medicine, University of Pennsylvania, Philadelphia, PA, USA; Department of Biomedical Sciences, School of Veterinary Medicine, University of Pennsylvania, Philadelphia, PA, USA; Department of Biology, University of Pennsylvania, Philadelphia, PA, USA; Department of Pathology & Laboratory Medicine, Perelman School of Medicine, University of Pennsylvania, Philadelphia, PA, USA; Abramson Family Cancer Research Institute, Perelman School of Medicine, University of Pennsylvania, Philadelphia, PA, USA; Department of Genome Sciences, University of Washington, Seattle, WA, USA; Department of Cell & Developmental Biology, Perelman School of Medicine, University of Pennsylvania, Philadelphia, PA, USA; Department of Molecular & Systems Biology, Dartmouth Geisel School of Medicine, Lebanon, NH, USA; Allen Discovery Center for Cell Lineage Tracing, Seattle, WA, USA; Brotman Baty Institute for Precision Medicine, Seattle, WA, USA; Howard Hughes Medical Institute, Seattle, WA, USA; Institute for Regenerative Medicine, University of Pennsylvania, Philadelphia, PA, USA

## Abstract

Metastatic cancer remains largely incurable due to an incomplete understanding of how cancer cells disseminate throughout the body. However, tools for probing metastatic dissemination and associated molecular changes at high resolution are lacking. Here we present multiplexed, activatable, clonal, and subclonal GESTALT (macsGESTALT), an inducible lineage recorder with concurrent single cell readout of transcriptional and phylogenetic information. By integrating multiple copies of combined static barcodes and evolving CRISPR/Cas9 barcodes, macsGESTALT enables clonal tracing and subclonal phylogenetic reconstruction, respectively. High barcode editing and recovery rates produce deep lineage reconstructions, densely annotated with transcriptomic information. Applying macsGESTALT to a mouse model of metastatic pancreatic cancer, we reconstruct dissemination of tens-of-thousands of single cancer cells representing 95 clones and over 6,000 unique subclones across multiple distant sites, e.g. liver and lung metastases. Transcriptionally, cells exist along a continuum of epithelial-to-mesenchymal transition (EMT) *in vivo* with graded changes in associated signaling, metabolic, and regulatory processes. Lineage analysis reveals that from a majority of non-metastatic, highly epithelial clones, a single dominant clone that has progressed along EMT drives the majority of metastasis. Within this dominant clone a parallel process occurs, where a small number of aggressive subclones drive clonal outgrowth. By precisely mapping subclones along the EMT continuum, we find that size and dissemination gradually increase, peaking at late-hybrid EMT states but precipitously falling once subclones are highly mesenchymal. Late-hybrid EMT states are selected from a predominately epithelial ancestral pool, enabling rapid metastasis but also forcing extensive and continuous population bottlenecking. Notably, late-hybrid gene signatures are associated with decreased survival in human pancreatic cancer, while epithelial, early-hybrid, and highly mesenchymal states are not. Our findings illuminate features of metastasis and EMT with the potential for therapeutic exploitation. Ultimately, macsGESTALT provides a powerful, accessible tool for probing cancer and stem cell biology *in vivo*.

## Introduction

The vast majority of cancer deaths are due to metastasis, a process that transforms a localized, often curable lesion into a systemic, largely incurable disease^1,2^. Recurrent genetic drivers of metastasis have proven elusive, suggesting that other levels of dysregulation may principally drive the phenomenon^1^. Phylogenetic histories of cancer progression in individual patients, e.g. based on analyses of copy number variation (CNV) or somatic mutation, can inform how the cells comprising metastases are related to the primary tumor, as well as to one another^3^. However, such methods are restricted to natural genetic diversity and additionally fail to concomitantly capture the molecular phenotype of each profiled cell, limiting what can be learned about the cellular programs that underlie the development and relative success of metastases.

Beginning with GESTALT^4^ (<u>g</u>enome <u>e</u>diting of <u>s</u>ynthetic target <u>a</u>rrays for lineage tracing), a new paradigm for lineage tracing emerged, employing CRISPR/Cas9 to progressively and stochastically mutagenize a compact, genomically-integrated barcode, thereby producing patterns of edits that can be used to reconstruct phylogenetic relationships amongst cells^5^. Such methods can be readily coupled to single-cell RNA sequencing (scRNA-seq) to explicitly relate cell lineage histories with transcriptional states^6–8^. Until recently^9,10^, GESTALT and related methods have primarily been applied to early development, e.g. by injection of CRISPR/Cas9 reagents into zygotes and subsequent profiling of edited barcodes and single cell transcriptomes from the resulting multicellular organism. However, with refinement, CRISPR/Cas9-based lineage tracers have strong potential to be useful in other contexts, such as the study of somatic stem cell dynamics or cancer metastasis.

### An inducible lineage recorder with scRNA-seq readout

To this end, we developed macsGESTALT (<u>m</u>ultiplexed, <u>a</u>ctivatable, <u>c</u>lonal and <u>s</u>ubclonal GESTALT), an integrated, inducible, and scalable method that can be easily adapted to any engineerable mammalian system to enable lineage tracing (**Fig. 1a**). Our approach consists of three components: 1) Each cell contains multiple unique barcode integrations. Barcodes are constitutively expressed within the 3’ untranslated region (UTR) of a polyadenylated puromycin transcript, enabling sequencing via standard mRNA-based capture.

**Figure 1:**
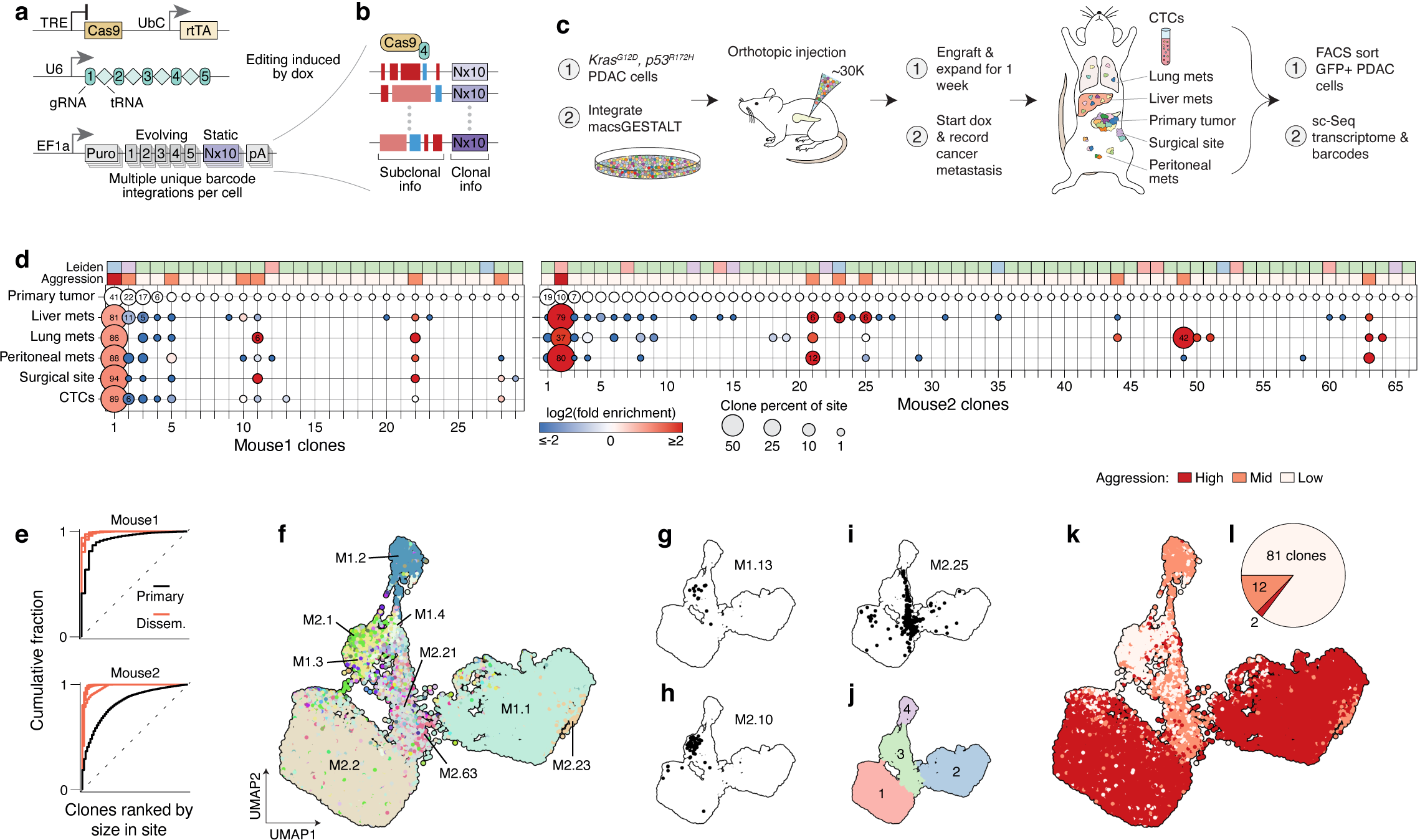
Most metastases arise from rare, transcriptionally-distinct clones. **a**, Genetic components of macsGESTALT, a broadly-applicable, inducible, and high-resolution lineage tracing system. **b**, Clone-level information is stored in static barcodes, while subclonal or phylogenetic information is dynamically encoded into evolving barcodes via indels (red and blue bars) induced by administration of doxycycline. **c**, Schematic of procedure for single-cell transcriptional profiling and lineage reconstruction of metastatic pancreatic cancer across 6 diverse harvest sites, including circulating tumor cells (CTCs). **d**, 95 clones across 2 mice reconstructed by static barcodes. Clones are numbered by size in the primary tumor (largest to smallest). Percent contribution to each harvest site (circle size) and enrichment compared to the primary tumor (circle color) are visualized. Top annotations show each clone’s Leiden transcriptional cluster and aggression assignments as in **j** and **k**, respectively. **e**, Cumulative fraction of each clone in each disseminated site (red) and primary tumor (black). Dotted-lines represent perfect clone-size equality. **f**, UMAP plot of 28,028 single cells containing both lineage and transcriptional information. Cells are colored by clone, with select large clones highlighted (as mouse.clone). **g, h**, Two representative non-aggressive clones occupying similar transcriptional space. **i**, A representative clone of medium aggression. **j**, Leiden transcriptional clustering of **f. k**, Cells in UMAP space annotated by clonal aggression. **l**, Number of non-, mid-, or high-aggression clones of 95 total.

Each barcode is a combination of a static 10bp sequence of random bases, used for clonal reconstruction, and a 250bp editable, evolving region composed of five CRISPR target sites, used for phylogenetic reconstruction (**Fig. 1b**). 2) The evolving region is targeted by an array of five guide RNAs (gRNAs), separated by transfer RNA (tRNA) spacers, under a single constitutive mammalian U6 promoter. Upon transcription, tRNAs are excised from the array by endogenous RNAse P and Z, releasing the individual gRNAs^11^. We selected this configuration from a screen of five different arrays, as it was compact and could easily be placed under different promoters as needed, yet drove robust barcode editing (**Extended Data Fig. 1**). 3) Cas9 expression and barcode editing are induced by doxycycline (dox) binding to a constitutive reverse tetracycline transactivator (rtTA) and activating a tetracycline responsive element (TRE) promoter^12^. Inducible barcode editing *in vitro* was robustly driven with limited leakiness, mostly confined to the first target site (**Extended Data Fig. 2, 3**). We also validated successful barcode recovery and clonal reconstruction in two independent experiments, each involving limiting dilution, expansion, and single cell sequencing (**Extended Data Fig. 4**).

### Aggressive clones are rare and transcriptionally divergent

We next set out to investigate cancer metastasis at high resolution by combining macsGESTALT and scRNA-seq^6,8^. We focused on pancreatic ductal adenocarcinoma (PDAC), which has a 5-year survival rate of 9%, the lowest of any major cancer^13^. Furthermore, 90% of PDAC patients have some dissemination at the time of diagnosis^13^. To study PDAC metastasis, we employed a commonly used model, where cells from KPCY (*LSL-Kras*^*G12D/+*^; *Trp53*^*LSL-R172H/+*^; *Pdx1-cre; LSL-Rosa26*^*YFP/YFP*^) mouse tumors^14–16^ are orthotopically transplanted into the pancreata of non-tumor-bearing mice^15,17^. This approach faithfully models human disease, due to the following: 1) *Kras* gain-of-function and *p53* loss-of-function are the most common drivers of human PDAC^18^; 2) cells experience minimal time *in vitro* — a drawback of traditional cell lines; 3) a focal lesion develops in the pancreas that 4) disseminates to the same sites as human PDAC, including the liver and lung.

To investigate PDAC metastasis and associated transcriptional states, we selected a highly metastatic line from a library of characterized PDAC lines previously-derived from KPCY tumors^16^ (**Methods: Cell lines**). To enable lineage tracing of these cells, we introduced dox-inducible Cas9 and the gRNA array through lentiviral transduction, and separately introduced multiplexed barcodes via PiggyBac-transposition, thereby producing macsGESTALT PDAC cells. To model cancer metastasis *in vivo*, we injected mouse pancreata with 30,000 macsGESTALT PDAC cells, representing thousands of static barcode clones (**Fig. 1c**). After one week of engraftment, we administered doxycycline in the drinking water to begin lineage tracing. As expected^17^, all mice were morbid at five weeks post-injection. We randomly selected two mice, M1 and M2, and harvested cells from six cancer-bearing sites (primary tumor, liver, lung, peritoneal mets, surgical-site met, circulating tumor cells). PDAC cells were fluorescence sorted and processed for scRNA-seq of transcriptomes and macsGESTALT barcodes.

Overall, 89% of transcriptomes had corresponding clonal lineage information for M1 and 77% for M2, demonstrating improved barcode recovery using macsGESTALT compared to prior methods^6,9^. In total, across all sites in both mice, we recovered both the transcriptome and clonal history for 28,028 single cells (M1: 12,657; M2: 15,371) (**Extended Data Fig. 5a**,**b**). The set of static barcodes defining a clone were determined via hierarchical clustering and custom pipelines (**Methods: Clonal reconstruction and multiplet elimination**). Cells were then sorted into each clone based on their static barcode content, permitting even cells with missing barcodes to be assigned to the appropriate clone, while enabling explicit multiplet detection and filtration and resulting in only ∼0.5% unmatched cells (M1: 0.54% and M2: 0.51%) (**Extended Data Fig. 5a**). For M1, an average of 3.7 out of a possible 5.9 barcodes were recovered per cell, while recovery for M2 was 1.7 of 2.5 (**Extended Data Fig. 5a**), with the lower number of barcodes per cell in M2 likely contributing to its lower overall clonal lineage recovery.

Clonal reconstruction revealed 95 distinct clones across the two mice (**Fig. 1d**), identified by 227 static barcodes (**Extended Data Fig. 5a**), indicating that less than 1% of all injected clones successfully engraft. By contrast, *in vitro* experiments using the same cells and a similar time course revealed that most cells (clones) survive and form colonies on plates (**Extended Data Fig. 4**). Thus, cancer cells in this model experience dramatic bottlenecking during *in vivo* engraftment.

Among the surviving clones, fitness differences were pronounced and shaped population structure across sites (**Fig. 1d**,**e**). In the primary tumor, the majority (>50%) of cells came from a minority of clones (2 clones in M1; 6 clones in M2). Bottlenecking was even more extensive at metastatic sites, wherein 80-90% of cells typically came from a single clone (**Fig. 1d**,**e**), and both mice had one clearly dominant clone across all disseminated sites (M1.1, M2.2). On the other hand, 51% of clones (48/95) failed to metastasize at all, suggesting that mutations in *Kras* and *p53* alone do not ensure metastatic success.

We next asked whether clones were transcriptionally distinct. Indeed, cells from the same clone clustered together in UMAP space (**Fig. 1f**). This was true of both large and small clones (**Fig. 1g-i**). Importantly, this finding extended to cells harvested from different sites, suggesting that cells retain their clonal transcriptional identity even after dissemination (**Extended Data Fig. 6**). These stable transcriptional differences may result from either epigenetic drift or large-scale copy number changes, the latter observed in our data (**Extended Data Fig. 7**) and a hallmark of PDAC chromosomal instability^19^.

### An EMT continuum associated with aggression

We next asked if differences in clonal behavior corresponded to transcriptional differences. While clones had distinct transcriptional identities, we found that many overlapped in UMAP space (**Fig. 1f-i**). Furthermore, 81% of clones (77/95 across both mice) primarily resided in a single transcriptional cluster, Cluster 3 (**Fig. 1d**,**j**). To relate transcriptional state to tumor aggression, we derived a clonal aggression scoring system based on clone size and dissemination (**Fig. 1d, Methods: Clonal reconstruction and multiplet elimination**). We found that 85% (81/95) of clones were non-aggressive and were transcriptionally similar, residing in a small region of the aforementioned Cluster 3 (**Fig. 1k**,**l**). Conversely, highly-aggressive clones were exceedingly rare, yet were a dominant contributor to transcriptional diversity, as illustrated by their large spatial distribution in the UMAP embedding (**Fig. 1k**).

We then sought to understand the specific transcriptional programs associated with aggression. While both mice were strikingly similar in terms of clonal composition (**Fig. 1d**), we focused on M1, since we harvested cells from more sites and recovered over twice as many barcodes per cell, which permits effective downstream subclonal reconstruction (**Extended Data Fig. 5a**,**b**). Reanalyzing the data apart from M2, non-aggressive clones again appeared transcriptionally similar to one another (**Fig. 2a**). Interestingly, these clones were enriched for expression of canonical epithelial markers, such as *Epcam, Muc1*, and *Cdh1* (**Fig. 2b-d, Extended Data Fig. 8a**). Conversely, mesenchymal markers, such as *Sparc, Zeb2*, and *Col3a1*, were enriched in cells of the aggressive clone, M1.1 (**Fig. 2e-g, Extended Data Fig. 8b**). Loss of epithelial genes and gain of mesenchymal genes are defining hallmarks of epithelial-to-mesenchymal transition (EMT)^20,21^.

**Figure 2:**
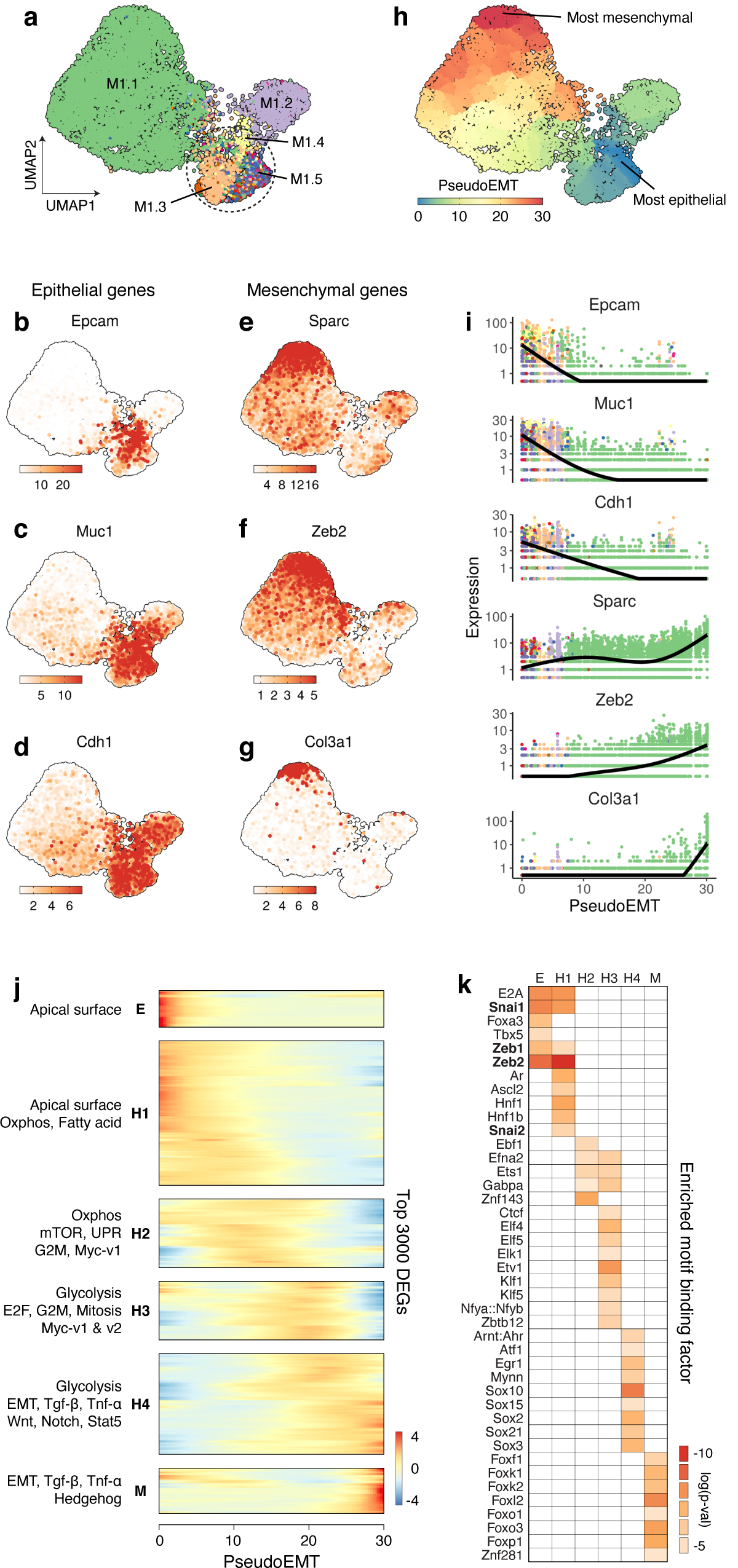
A transcriptional EMT continuum *in vivo*. **a**, UMAP plot of mouse 1 single cells, colored by clone, with the first 5 clones annotated. Circled region indicates the transcriptional space where smaller, non-aggressive clones reside. **b-g**, Expression of canonical epithelial (**b**,**c**,**d**) and mesenchymal (**e**,**f**,**g**) markers peaks on opposite ends of UMAP space. **h**, Unbiased trajectory inference revealing a pseudotime axis matching our EMT gene expression gradient (pseudoEMT). The root of the trajectory, or state of 0 pseudoEMT, is set to the region with most epithelial gene expression. **i**, Expression of EMT markers from **b-g** plotted along pseudoEMT and colored as in **a. j**, Hierarchical clustering of kinetic curves for the top 3000 differentially expressed genes across pseudoEMT (q = 0, Moran’s I > 0.1) for all cells from **h**. Gene clusters are labeled as epithelial (E), hybrid (H1, H2, H3, H4), or mesenchymal (M) based on their expression across pseudoEMT. Geneset analysis using MSigDB Hallmarks for each gene cluster (hypergeometric test, p < 0.05) **k**, Significantly enriched motifs (hypergeometric test, p < 0.05) in promoters for each gene cluster, with canonical EMT master regulators highlighted.

EMT is a process of transdifferentiation, wherein epithelial cells lose the properties of cell polarity and adhesion, while gaining the ability to be motile and migratory^20,21^. In cancer, EMT is implicated in invasion, metastasis, tumor stemness, plasticity, and drug resistance^20,21^. EMT is primarily a transcriptional process mediated by a group of key master-regulator transcription factors (EMT-TFs)^22^. We observed elevated expression in aggressive clones of 4/5 EMT-TFs, namely *Zeb1, Zeb2, Snai1*, and *Snai2* (**Fig. 2f, Extended Data Fig. 8c**). Expression of *Prrx1*, an important regulator of EMT in PDAC^23^, was also increased.

Traditionally, EMT was considered a binary process, where cells switch from fully epithelial to fully mesenchymal. However, recent studies have reported discrete intermediate EMT states^24–28^ or even a continuum of states^29,30^. In our data, epithelial and mesenchymal UMAP regions were not well segregated. Specifically, epithelial and mesenchymal genes appeared to gradually lose and gain expression as a function of distance from two extremes (**Fig. 2b-g**), suggesting that a continuum of EMT states exists *in vivo*.

We leveraged our single-cell data to explore the transcriptional correlates of EMT as a continuum. We performed unbiased trajectory inference using Monocle 3^31^ and found that the main trajectory in our data corresponded to the observed EMT gene expression axis (**Fig. 2h**). We named this trajectory “pseudoEMT” (akin to pseudotime for developmental trajectories) and placed the root of the trajectory, or the zero EMT state, at the most epithelial transcriptional region (**Fig. 2h**). Hence, the expression of canonical epithelial markers was highest at the root. We found that many genes, including known epithelial or mesenchymal markers, rise and fall at different rates across pseudoEMT (**Fig. 2i, Extended Data Fig. 9a-c**); for example, many extracellular matrix genes activate only very late in the trajectory (**Fig. 2i, Extended Data Fig. 9b**). Additionally, numerous genes, such as *Cd44* or *Inhba*, displayed unusual patterns, rising and then falling or plateauing (**Extended Data Fig. 9d**). Plotting cells along pseudoEMT also highlighted that smaller, non-aggressive clones reside on the epithelial extreme, while more mesenchymal states are restricted to large, aggressive clones, such as M1.2 and particularly M1.1 (**Fig. 2i**).

To systematically characterize gene expression along EMT, we identified the top 3000 significantly differentially expressed genes across pseudoEMT (q ∼ 0, Moran’s I > 0.1) (**Supplementary Table 1**). Hierarchical clustering of genes revealed six gene sets with similar kinetics (**Fig. 2j**). We classified these sets from most epithelial to most mesenchymal as follows: Epithelial (E), Hybrid 1, 2, 3, and 4 (H1, H2, H3, H4), and Mesenchymal (M) (**Fig. 2j, Supplementary Table 1**). We performed hypergeometric gene set enrichment using the Molecular Signatures Database (MSigDB) Hallmark gene sets, which represent well-defined biological states and processes (**Fig. 2j, Supplementary Table 2**). In concordance with the pseudoEMT trajectory, gene set enrichment indicated an EMT process. Early clusters (E, H1) were enriched for apical surface genes, consistent with epithelial cell polarity, while late clusters showed gradually increased enrichment for EMT (H4: p = 3×10^−6^, M: p = 3×10^−29^). An inducer of EMT and metastasis, TGF-β signaling^21,32,33^, as well as Jak/Stat3 and Stat5 signaling^34^, peaked in the late hybrid state (H4) and tapered off in the highly mesenchymal state (M). Other pathways purported to be involved in EMT, such as TNF-α^35^, Wnt^36,37^, and Hedgehog^38^ were also only enriched in H4 or M. Interestingly, Notch signaling was recently implicated as a hybrid-EMT stabilizer^39,40^, consistent with our finding that it was only enriched in H4.

Striking metabolic changes across EMT were also apparent. Transitioning from early (H1, H2) to late (H3, H4) hybrid gene clusters, we observed a strong shift from enrichment of oxidative phosphorylation (OXPHOS) toward glycolysis, potentially related to the enrichment of mTOR signaling in H2^41^. Consistent with metabolic shifts, hybrid EMT states also were highly enriched for proliferative gene sets, such as G2M, E2F, and mitotic spindle. Specifically, enrichment began modestly in H2 and peaked dramatically in H3 (G2M, H2: p = 3×10^−2^, H3: p = 1×10^−20^). We next determined the cell cycle phase of each cell (G1, G2M, or S) to estimate the proportion of actively dividing cells (S/G2M) across pseudoEMT (**Methods: Single cell transcriptome data processing**). Consistent with Hallmark gene set enrichment, cell cycling peaked at EMT regions representing the E and H2/H3 gene clusters (**Extended Data Fig. 9e**). These hybrid EMT proliferative changes were potentially driven by Myc^42^, as Myc targets mirrored proliferative gene set enrichment and cell cycling fraction (Myc-v1, H2: p = 1×10^−3^, H3: p = 1×10^−30^).

We next asked which TFs might regulate progression through EMT. Using HOMER^43^, we detected 45 significantly enriched DNA motif binding factors across all gene clusters (**Fig. 2k**). EMT master regulators, *Zeb1, Zeb2, Snai1*, and *Snai2*, were enriched in early clusters, E and H1. As EMT-TFs are primarily transcriptional repressors that downregulate epithelial genes^22^, this finding illustrates our ability to discover regulators of the EMT continuum. ETS-domain TFs, which are associated with metastasis, invasion, and EMT^44,45^, dominated the enrichment profiles of hybrid states H2 and H3. Motifs bound by members of the Sox and Fox families were enriched in H4 and M, respectively. Sox TFs are often associated with stemness-related processes^46^. Notably, the six gene clusters have no overlapping genes, yet adjacent clusters often displayed overlapping TF and gene set enrichment, lending further support for a gradual continuum of EMT transitions (**Fig. 2j**,**k**). Overall, across this continuum of 3000 genes, we describe many classic EMT markers, pathways, and regulators, but we also find many novel or less well-characterized genes and processes of potential interest for furthering understanding of EMT *in vivo* (**Supplementary Table 1, 2**).

### Late-hybrid EMT states are proliferatively and metastatically advantageous

Most cells in the mid-to-late EMT continuum came from a single dominant clone, M1.1, providing only coarse resolution into the transcriptional processes associated with tumor aggression and highlighting the limitations of static barcoding (**Fig. 2i**). We therefore leveraged editing patterns of macGESTALT evolving barcodes to more precisely relate EMT and aggression at the subclonal level.

We recovered a large number of edited and informative target sites per cell, conducive to phylogenetic analysis. Altogether, we observed an editing rate of 96% across 384,870 recovered target sites (**Extended Data Fig. 10a**). Editing was distributed across the length of the barcodes with peaks at the expected Cas9 cut-sites, 3bp upstream of the protospacer adjacent motifs (PAMs) (**Fig. 3a**). Deletions predominated over insertions as expected^4,6,9^, with an approximately equal number of single- and multi-target deletions (**Fig. 3b, Extended Data Fig. 10b**). The average edit size varied by edit type, with 11bp for insertions, 18bp for single-target deletions, and 80bp for multi-target deletions (**Extended Data Fig. 10c**). Multi-target deletions were of a large size range and involved 2, 3, 4, or 5 target sites at frequencies ranging from 10-19% (**Extended Data Fig. 10b**,**c**). Individual target site editing rates varied between 89-99% (**Fig. 3b**). We recovered an average of 18.5 target sites (3.7 barcodes) per cell for M1 and 8.5 (1.7) for M2 (**Extended Data Fig. 5a**).

**Figure 3:**
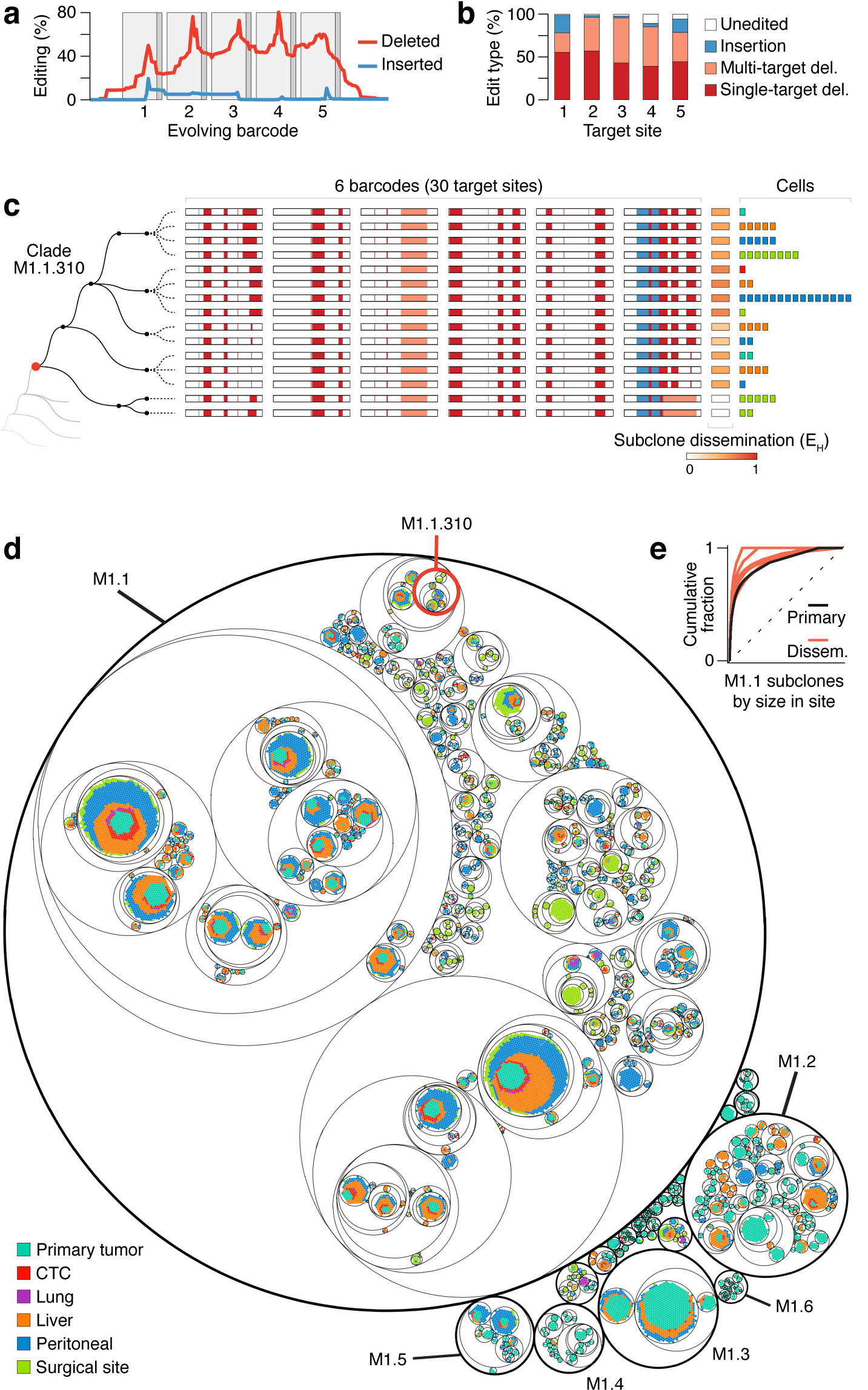
High-resolution subclonal lineage reconstruction of metastatic cancer. **a**, Percent at which each base is mutated along the length of 227 evolving barcodes from 95 clones across both mice. Target site spacers (light grey) and PAMs (dark grey) are indicated. **b**, Edit types observed at each target site across the same evolving barcodes as in **a. c**, Example phylogenetic reconstruction of a small clade within clone M1.1. Clade M1.1.310 (root node in red) contains 6 distinct subclones composed of 58 cells from 5 different harvest sites. Each cell in this clade has 6 evolving barcodes, illustrated by white bars with colored edits overlaid. Cells with the same barcode editing pattern are grouped into a subclone; subclone phylogeny is inferred from common and distinct edits. Dotted lines emerging from each subclone node represent the harvest sites from which cells were recovered. Subclone dissemination is a statistical metric of how equally a subclone’s cells are distributed across sites (Shannon Equitability, adjusted for sampling size), where 0 is no dissemination. Individual cells are stacked and colored by site on the far right. **d**, Circle packing plot of the full single cell phylogeny of mouse 1, with clade M1.1.310 from **c** circled in red. Outermost circles (heavy black borders) define individual clones, with the 6 largest clones labeled. Within each clone, nested circles group increasingly related cells based on their barcode editing patterns. Innermost circles contain cells from reconstructed subclones. Each point represents a single cell, colored by harvest site. **e**, Cumulative fraction of each subclone of clone M1.1 in each harvest site. Dotted-line represents perfect subclone-size equality.

Intraclonal tree reconstruction was performed in three main steps (**Fig. 3c**). First, different barcodes from the same cell were concatenated based on their static barcodes into a “barcode-of-barcodes”, which contains all of the phylogenetic information recovered for that cell. Second, cells with identically edited barcode-of-barcodes were grouped into subclones, since they are indistinguishably close relatives. Third, phylogenetic relationships between subclones were reconstructed based on edit inheritance patterns (**Fig. 3c**). Subclonal metastatic aggression was quantified via Shannon’s Equitability (E_H_) – a statistical measure of dissemination across harvest sites (**Methods: Subclonal dissemination calculation**). For example, a subclone found at only one harvest site is not metastatically aggressive and has an E_H_ of zero.

We recovered 6,055 unique barcode-of-barcodes, which yielded 1,692 transcriptionally-useful subclones across 95 clones (**Extended Data Fig. 10a, Methods: Subclonal and phylogenetic reconstruction**). 903 of these subclones were from M1, as it contained over twice as many edited target sites per cell compared to M2 (**Extended Data Fig. 10a**). The greater reconstructive power for M1 was particularly apparent in the difference between the critical dominant clones of each mouse, where M1.1 with seven barcode integrants had 601 subclones compared to M2.2 with only two integrants and 110 resulting subclones. The full clonal and subclonal phylogenetic visualization of M1 highlights the overwhelming proliferative and metastatic dominance of clone M1.1 (**Fig. 3d**). However, within M1.1, we also observed vast heterogeneity with respect to subclonal aggression and metastatic success. Most strikingly, the same bottlenecking observed on the clonal level was also present on the subclonal level within M1.1 (**Fig. 3e**). Subclonal bottlenecking further increased at metastatic sites, again mirroring observations at the clonal level. Thus, cancer progression appears to be defined by a state of constant competition and selection, separate from the effects of engraftment.

To understand how differences in subclonal behavior may relate to EMT, we calculated the mean pseudoEMT value for each subclone and plotted this and subclonal dissemination (E_H_) for clone M1.1 (**Fig. 4a**,**b**). While M1.1 was highly mesenchymal compared to other M1 clones, many subclones within M1.1 were actually quite epithelial. These epithelial subclones were primarily small and non-metastatic (**Fig. 4a**,**b**). Interestingly, the same was true of highly mesenchymal subclones. On the other hand, the largest and most disseminated subclones appeared to express hybrid EMT states (**Fig. 4a**,**b**), supporting the idea that EMT extremes are less metastatic than hybrid states^21,28,47,48^.

**Figure 4:**
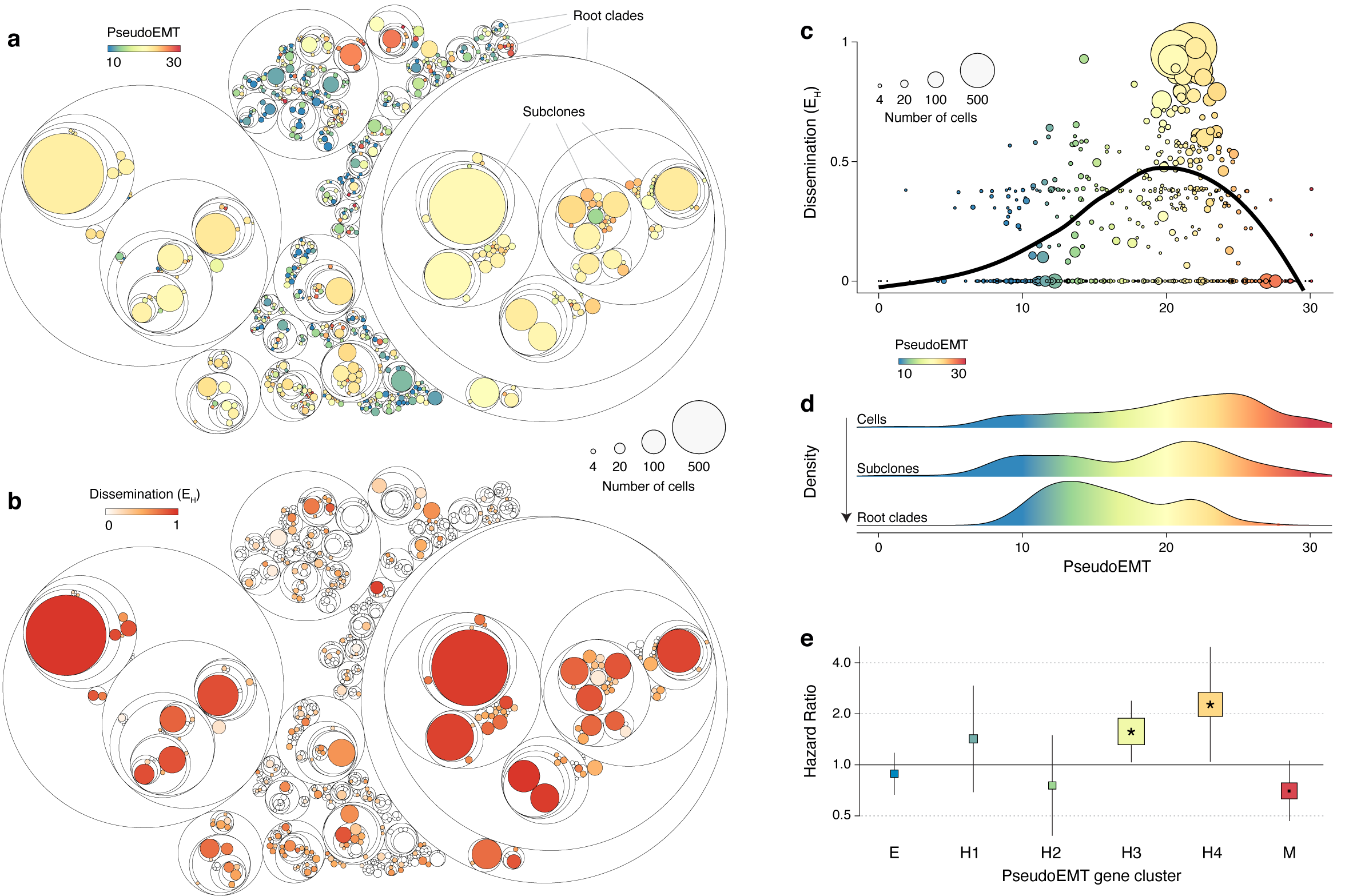
Peak metastatic aggression corresponds to late hybrid-EMT transcriptional states. **a, b**, Circle packing plots of the phylogenetic structure of clone M1.1 with subclones colored by mean pseudoEMT (**a**) and by dissemination score (**b**). **c**, Relationship between metastatic dissemination and pseudoEMT transcriptional state for all subclones from **a** and **b. d**, Density along pseudoEMT of M1.1 cells and their increasingly ancestral (arrow) phylogenetic groups, i.e. subclones and root clades (examples highlighted in **a**), illustrating that epithelial transcriptional states are outcompeted over time. **e**, Relationship between PDAC patient survival (TCGA-PAAD, n=173) and patient enrichment scores for each pseudoEMT gene cluster (E, H1, H2, H3, H4, M) using Cox regression analysis, with the hazard ratio for each gene cluster displayed (95% CI, *, p<0.05, ·, p<0.1). Square sizes are inversely proportional to p-value.

To precisely characterize where aggression peaked along the EMT continuum, we mapped subclonal dissemination (E_H_) and size along pseudoEMT (**Fig. 4c**). We found that dissemination gradually peaked around the H3 and H4 hybrid states (pseudoEMT score of 20-22) and then sharply declined at highly mesenchymal states. Thus, late-hybrid EMT states are metastatically advantageous and are associated with specific proliferative, metabolic, and signaling processes (**Fig. 2j**), as well as distinct regulatory binding factors (**Fig. 2k**).

Notably, hybrid-EMT states appeared transcriptionally stable – for example, a large, hybrid subclone often had close relatives that were also large and hybrid (**Fig. 4a**). To understand the stability of EMT states, we plotted the distribution of cells, subclones, and root clades along pseudoEMT (**Fig. 4d, Methods: PseudoEMT across ancestral relationships**). Root clades mark the first phylogenetic subdivision within a clone and are hence an older subgrouping of cells than a subclone. Examples of root clades and subclones are highlighted in **Fig. 4a**. Root clades exist at the time of dox initiation (one week post orthotopic transplant), cells exist at the time of harvest, and subclones in between; thereby we compared different levels of ancestral groups. Moving from root clades to cells, there was a shift from epithelial to hybrid states, suggesting that while epithelial states are the prevailing default, they are outcompeted by hybrid states (**Fig. 4d**). This intraclonal observation again mirrored findings at the clonal level, where most clones were epithelial but were heavily outcompeted by the dominant clone M1.1. Therefore, ongoing natural selection of rare, late-hybrid EMT states over predominating epithelial states permits both rapid dissemination and forces continuous clonal and subclonal bottlenecking.

As late-hybrid EMT states, namely the H3 and H4 gene clusters, were profoundly associated with metastasis in our model, we asked whether a similar trend might exist in human PDAC (**Fig. 4e**). Using The Cancer Genome Atlas (TCGA) matched gene expression and clinical data, we found that while the E, H1, and H2 gene clusters had no association with disease prognosis, patients enriched for H3 and H4 had a significantly increased risk of death and that this risk disappeared for the highly mesenchymal cluster M (**Fig. 4e**). Remarkably, these human PDAC findings faithfully mirror the rise and fall of subclonal metastatic aggression along pseudoEMT in our model (**Fig. 4c**).

## Discussion

Recurrent genetic drivers of metastasis remain lacking, suggesting that non-genetic adaptations may be key to understanding cancer dissemination. To study such processes at high-resolution, we developed macsGESTALT, a flexible, inducible lineage tracer that can be easily coupled with scRNA-seq. In applying macsGESTALT to understand pancreatic cancer metastasis, we gained critical insight into cancer behavior at the clonal, subclonal, and transcriptomic levels. Despite using a metastatically competent genetic model, we found that most clones do not metastasize, supporting the importance of transcriptional and non-genetic processes in metastasis^1^. While non-aggressive clones occupy similar transcriptional space, many aggressive clones conversely have distinct transcriptional identities, which they retain even upon dissemination to distant metastatic sites. Among aggressive clones, we found that a single dominant clone drives the overwhelming majority of metastasis across all sites, without apparent organotropism. As these findings are remarkably consistent across both mice — which faithfully recapitulated the kinetics of cancer morbidity following orthotopic PDAC transplant^17^ — we suspect the emergence of rare dominant clones from many non-metastatic clones may be a conserved feature of metastasis in this model.

Highlighting the limitations of static barcoding approaches in isolation, extensive clonal bottlenecking obscured lineage information at critical points *in vivo*. To address such challenges, macsGESTALT pairs static barcodes for clonal reconstruction with inducible evolving barcodes for subclonal reconstruction. In this study, evolving barcodes revealed that growth and dissemination of the dominant clone are driven by rare highly aggressive subclones that associate with specific EMT transcriptional states. While a wide-range of EMT states exist — from highly epithelial to highly mesenchymal — aggressive subclones exhibit primarily late-hybrid EMT states. These late-hybrid subclones appear to undergo continuous and aggressive evolutionary selection from a background of predominantly epithelial states. While this process enables rapid proliferation and metastasis, it necessitates extensive population bottlenecking, presenting a potentially vulnerable or exploitable feature of PDAC metastasis. Further highlighting the therapeutic relevance of these findings, late-hybrid EMT states correspond with worse overall survival in human PDAC, while epithelial, early-hybrid, or highly mesenchymal states do not, thereby mirroring the rise and fall of metastatic capability across pseudoEMT in our model. As such, we characterized the EMT spectrum in depth, finding numerous enriched signaling, metabolic, and regulatory features throughout. Amongst these, in late-hybrid EMT states, we observed increased MYC activity and proliferation, as well as potential metabolic rewiring from OXPHOS to glycolysis, which has been implicated in both tumor invasiveness^49,50^ and EMT^51,52^.

By exploring the dynamics of a dominant clone driving metastasis in a mouse model of PDAC, we characterized a detailed molecular roadmap of EMT *in vivo* and highlight one potential path to aggressive metastasis. As cancers are notoriously heterogeneous, we anticipate that many different paths to aggressive dissemination likely exist — but promisingly, we find that the late-hybrid EMT states uncovered here also predict worse survival in a large human patient cohort, suggesting they may be a conserved mechanism. As PDAC has the lowest survival rate of any major cancer^13^, largely due to aggressive, early metastasis present at diagnosis, we hope that our approach will enable future studies to reveal additional processes underlying the highly metastatic nature of PDAC.

Our insights derive from a global, unbiased assessment of metastatic phylogeny and transcription at the single cell level. macsGESTALT enables such investigations by combining static and evolving lineage tracing and achieving high barcode recovery and editing rates, producing rich lineage trees densely annotated with transcriptional information. Furthermore, the inducibility of macsGESTALT allows lineage tracing to initiate at the optimal experimental time, here after tumor engraftment. However, a future application could be to couple initiation of macsGESTALT with a specific intervention to study its effects on population structure, such as administration of a therapy or an injury model. As macsGESTALT only requires integration of three components and is readily coupled with single-cell sequencing, we hope that it will be rapidly adopted to address questions in cancer and stem cell biology at previously inaccessible levels of resolution and scale.

## Methods

### Plasmid design and construction

All Gibson assemblies were performed using NEBuilder HiFi DNA Assembly Master Mix (NEB #E2621) and were assembled at 50 °C for 60 min at appropriate molar ratios. For cloning, all PCRs were performed using HotStart ReadyMix (Kapa Biosystems #KK2601). Restriction enzymes, instead of PCR, were used to linearize vector backbones to prevent backbone mutations. All bacterial transformations were performed with NEB Stable Competent E. coli (NEB #3040H) and cells were grown at 30 °C for 24 h, unless otherwise noted. Final plasmid preps were performed with Zymopure II Plasmid Kits (Zymo Research #D4202). All regulatory, coding, and editing-related regions in final assembly products were validated by Sanger sequencing. All gene block sequences were ordered from IDT. Full, annotated sequences are available via Benchling for all plasmids generated or used for editing experiments; macsGESTALT plasmids will be deposited in Addgene.

V7 and V8 barcoding lentiviral transfer plasmids used for guide RNA array screening were constructed in 2-part Gibson assemblies using pLJM1-EGFP (Addgene #19319)^53^ backbone digested with EcoRI + gene blocks for V7 or V8 barcodes to make pLJM1-EGFP-V7 and pLJM1-EGFP-V8.

pUltra-U6-crRNAs-U6-tracr was constructed in a 3-part Gibson assembly using PacI linearized pUltra (Addgene #24129)^54^ backbone, a U6-driven array of 10 V8 targeting crisprRNAs (crRNAs) interspersed by tRNAs ordered as a gene block (pUltra5-U6crRNA-GA1), and another gene block encoding a U6-driven tracrRNA (GA1-U6-tracr-pUltra3).

The dox-inducible crRNA array plasmid, pBS31-GFP-V8crRNAs-U6-tracr-Ub-M2rtTA, was constructed in a 3-part Gibson assembly using EcoRI linearized pBS31, a gene block containing 10 V8 targeting crRNAs interspersed by tRNAs in the 3’ of a GFP opening reading frame (ORF) (TP-gB-1), and a gene block containing U6-driven tracrRNA followed by Ubc promoter-driven M2-rtTA with a V8 barcode of 10 targets in the 3’ UTR (TP-gB-2). The barcode was excised for transient transfection gRNA screening experiments by digesting with NsiI and religating the backbone.

p5xU6_5sgRNA-Hsp70-Cas9GFP-pA that had V7 gRNAs 5-9 each with a separate U6 promoter was a gift from J. Gagnon^6^.

pCFDg1-5 gRNA-tRNA array was constructed stepwise as previously described using pCFD5 (Addgene #73914)^11^ as a template and V8 targeting gRNAs.

pUltra-U6-gRNAs1-5 lentiviral transfer plasmid, which was used to make macsGESTALT PDAC cells, was generated in a 3-part Gibson assembly using pUltra backbone linearized with PacI, a gene block with U6 promoter and gRNA 1 (pUltra5-U6-gRNA1), and a PCR-amplicon, amplified from pCFDg1-5, containing gRNA-tRNAs 2-5 (gRNAs1-5-pUltra3), thereby producing a constitutively-expressed five gRNA-tRNA array and a constitutive GFP selection marker.

PB-EF1α-Puro-V8.2 library cloning was performed as a 3-part Gibson assembly: 1) PB-CMV-MCS-EF1α-Puro (Systems Biosciences PB-510B-1) was digested with SpeI and HpaI to excise its cargo and create a linear backbone. 2) EF1α promoter and puro resistance gene were amplified from lentiGuide-Puro (Addgene #52963). 3) The V8.2 target array was ordered as a gene block. This assembly produced the PB-EF1α-Puro-V8.2 vector. Then, the barcode library was generated via a 2-part Gibson assembly using EcoRI linearized PB-EF1α-Puro-V8.2 and a random 10 bp containing staticID (static barcode) fragment, which was made by annealing and extending a pair of oligos (targetbarcode-r: TTTGTCCAATTATGCTCGAGGTCGAGAATTNNNNNNNNNNCGTTGATCGCACGCCA, targetbarcode-f2: TAGTTGGTTCCTACTGGCGTGCGATCAACG). The library was transformed into NEB 10-beta Electrocompetent E. coli (NEB #3020K), and the entire transformation was grown as a midi culture and prepped with Chargeswitch Pro Filter Midi Kit (Thermofisher #CS31104).

### Cell lines

Lentiviruses were packaged in HEK 293T cells using psPAX2 (Addgene #12260) and pMD2.G (Addgene #12259) second generation packaging and envelope plasmids. Viral supernatants were collected 2-4 d post-transfection and filtered through 0.45 µm filters. Filtered supernatants were either stored at -80 °C (never refrozen) or used fresh to infect cells. Barcoded 293T cells for the gRNA screen were produced by infecting with pLJM1-EGFP-V7 or pLJM1-EGFP-V8 lentivirus at low MOI (MOI < 0.2) and sorted by fluorescence-activated cell sorting (FACS) for GFP using a BD FACSAria II (BD Biosciences).

For the PDAC cells used to generate macsGESTALT PDAC cells, we selected the most metastically aggressive cell line (6419c5) from a published library of clonal PDAC lines^16^, which were each derived from harvested KPCY tumors. While this cell line originated from a single cell bottleneck during derivation, it had since been passaged ∼15x, thereby overtime in culture, becoming effectively polyclonal at the point of macsGESTALT barcode delivery.

macsGESTALT components were introduced into PDAC cells in 3 steps: First, dox-inducible Cas9 was integrated with Lenti-iCas9-neo (Addgene #22667)^12^, and infected cells were selected for neomycin resistance via G418 for 7 d. Second, the cells were infected with pUltra-U6-gRNAs1-5 at high MOI (MOI > 0.8), and the top 50% of GFP positive cells were sorted by FACS using a BD FACSAria II. This step was repeated once to produce cells with high gRNA array expression to ensure a high editing rate. This can be decreased to slow and spread the editing rate over time. Third, cells from the previous steps were barcoded by cotransfecting PB-EF1α-Puro-V8.2 library and Super PiggyBac Transposase plasmid (SBI #PB210PA-1) at a 1:10 molar ratio using Lipofectamine 3000 (Thermofisher). Barcoded cells were puromycin-selected for 7 d. To maintain diversity and limit leaky editing, cells were expanded after withdrawal of purmycin and frozen down with minimal time in culture (< 7 d). For lineage tracing experiments, cells were only expanded after thawing for 2-4 d as needed prior to orthotopic injection or experiment start.

### Guide RNA array editing screen

293T cells were cultured in culture media (DMEM, 10% FBS, 1% glutamine with penicillin and streptomycin). 293T cells barcoded with pLJM1-EGFP-V7 or pLJM1-EGFP-V8 lentivirus were transiently transfected with different combinations of plasmids to test gRNA array editing efficacy. Barcoded cells plated at 250,000 cells per well of 6-well plates, and transfected the following day with Lipofectamine 2000 (Thermofisher #11668030). 1.5 µg of px330 was used in each well (except no-transfection and pUltra-only control wells). All wells receiving a gRNA array plasmid were also transfected with a 1:1 molar amount of the appropriate gRNA plasmid compared to px330. Dox was initiated where appropriate the day after transfection. Additionally, as a positive control, one well received px330 and *in vitro* transcribed (IVT) gRNAs. Guide templates matching the V8 target sites were constructed and transcribed using GeneArt Precision gRNA Synthesis Kit (Thermofisher #A29377); gRNA 6 and 7 IVT reactions failed and these guides were excluded from further steps. IVT gRNAs were transfected using Lipofectamine CRISPRMax (Thermofisher #CMAX00001) 24 h after px330 was transfected. Expression of plasmids containing fluorescent markers was confirmed by microscopy. Cells were then allowed to expand and edit for one week and then harvested for library preparation and sequencing.

### PDAC dox-induced *in vitro* editing experiments

PDAC cells were cultured in complete media (DMEM, 10% FBS, 1% glutamine with penicillin and streptomycin). Dox-induced editing checks of macsGESTALT PDAC cells were performed in two separate experiments: In the first experiment, cells were plated and started on dox at 3 doses, 0, 0.1, or 2 µg/mL, with media change every other day. Cells were collected at 2 timepoints — after 1 and 2 weeks of dox exposure — and harvested for library preparation and sequencing. In the second experiment, cells were kept on 6 different dosages of dox, 0, 10, 50, 100, 500, or 1,000 ng/mL, for 2 weeks and harvested for library preparation and sequencing.

### Bulk genomic DNA barcode library prep, sequencing, and analysis

For all bulk DNA editing experiments, approximately one million cells were harvested per condition, washed, pelleted, and genomic DNA extracted with the NucleoSpin DNA RapidLyse Kit (Macherey-Nagel #740100.50). Genomic DNA was normalized to 30-50 ng/µL for each sample. All PCR reactions were performed using SYBR-containing master mix from the KAPA Real-Time Library Amplification Kit (Kapa Biosystems #KK2702) and terminated in the mid-exponential phase to limit over-amplification. AMPure beads (Agencourt Beads, Beckman Coulter #A63880) were used at a ratio of 1.5x to purify products after all PCR reactions. Barcodes were amplified from genomic DNA in a nested approach and sequencing adaptors, sample indices, and flow cell adaptors were added by a series of subsequent PCRs. For 293T samples containing pLJM1-EGFP-V7 or pLJM1-EGFP-V8, barcodes were amplified and adaptors added in a series of 3 PCRs. For PDAC samples containing PB-EF1α-Puro-V8.2, barcodes were amplified and adaptors added in a series of 4 PCRs. Primer sequence, purpose, and annealing temperature for all PCRs in both of these library preparations are included in **Supplementary Table 3**. In all cases, 250 ng of genomic DNA was loaded into a 50 µL PCR. Sample indices were added using NEBNext Multiplex Oligos for Illumina (Dual Index Primers Set – New England Biolabs). The concentration of final amplicons was measured by Qubit and the length validated by TapeStation HSD1000 prior to sequencing using Illumina MiSeq 600-cycle v3 Reagent Kits with the following run parameters: Read 1 - 301 cycles, i7 index - 8 cycles, i5 index - 8 cycles, Read 2 - 301 cycles. Bulk sequencing data for all samples was processed as previously reported^4^ and available on Github (https://github.com/mckennalab/SingleCellLineage/), with the UMI option set to FALSE (no UMI used). Output files were used for generating visualizations using the R programming language.

### Limiting dilution PDAC experiments and 10x Chromium loading

macsGESTALT PDAC cells were plated in a limiting dilution of ∼5 or ∼100 cells per well in a 48-well plate. Single cells gave rise to colonies and expanded. Cells were all allowed to expand without split for 2 weeks. The 100-cell wells were confluent and overgrown after 1 week in culture. The 5-cell wells were approximately 80-90% confluent at 2 weeks. At 2 weeks, a healthy, representative well from each condition was selected and passaged at a 1:2 split into a well of a 6-well plate. After 3 d, cells were harvested and dissociated using 500 µL TrypLE (Thermofisher #12605010) for 3-5 min. Reactions were neutralized with 3 mL culture media. Cell clumps were further dissociated by gently pipetting up and down 10x with a p1000, and then cells were centrifuged at 250*g* for 5 min. Cells were gently resuspended with a p1000 in 1 mL culture media, filtered through a 30 µm strainer, ensured to be in a single cell suspension under a light microscope, and counted with a hemocytometer. Cells were washed twice with 1 mL cold HBSS with 0.04% BSA (centrifuged at 150*g* for 3 min each time). Cells were filtered again through a 30 µm strainer and resuspended in cold HBSS with 0.04% BSA at a concentration of 700 cells/µL. Cells were counted again with a hemocytometer to ensure accurate concentration. For the 5-cell dilution sample, 8,000 cells were loaded on 10x (Chromium Single Cell 3’ Reagent Kits v3) targeting 5,000 cell recovery; for the 100-cell dilution sample, 16,000 cells were loaded targeting 10,000 cell recovery.

### Mice and orthotopic injection metastasis model

macsGESTALT PDAC cells were thawed and expanded for 2-4 d prior to dissociation and orthotopic injection into 10 week old NOD/SCID male mice (Jackson Laboratory). Approximately 30,000 PDAC cells were injected into the surgically-exposed tail of the pancreas, as previously described in detail^17^. Cells were allowed to engraft; then doxycycline was initiated 1 week post-injection and given continuously in the drinking water at 1 mg/mL. Mice were harvested at approximately 5 weeks post injection, once reaching morbidity. Primary tumor (PT), liver, lung, peritoneal macrometastases, and surgical-site lesions were sorted for both mice. Due to a more productive blood-draw, circulating tumor cells (CTCs) were captured for M1 but not M2. Additionally, the surgical-site lesion, which is similar in size and location to other peritoneal macrometastases, was processed separately in M1 but not M2. All mice were maintained in a specific pathogen-free environment at the University of Pennsylvania Animal Care Facilities. All experimental protocols were approved by and performed in accordance with the relevant guidelines and regulations of the Institutional Animal Care and Use Committee of the University of Pennsylvania.

### Blood harvest and preparation

When harvesting tissues, blood was extracted first via cardiac puncture using a 25 gauge 5/8 needle with 1 mL syringe attached. A successful blood draw was 400-700 µL, which was immediately transferred to a FACS tube containing 4% sodium-citrate in Milli-Q water. This was pelleted at 500 g for 5 min and red blood cells were lysed by resuspension in 2 mL ACK (Ammonium-Chloride-Potassium) buffer and incubation for 5 min at room temperature. 3 mL PBS were added and the mix was pelleted at 500 g for 5 min. Red blood cell lysis was repeated 2 times. Finally, cells were resuspended in 400 µL of cold FACS buffer (PBS, 2% FBS, 1 mM EDTA, 40 ug/mL DNase) with DAPI and strained through a 35 µm filter for FACS.

### Primary tumor, peritoneal macrometastases, and surgical-site harvest and dissociation

Primary tumor and macrometastases (metastases that could be manually handled, including surgical-site lesion) were excised from surrounding tissue, removing as much normal surrounding tissue as possible. All macrometastases from a mouse were processed as one sample. Samples were then transferred to a 6-well plate and washed with cold PBS 3x. Samples were minced, then transferred into 10 mL of DMEM containing 2 mg/mL collagenase IV plus 40 µg/mL DNase and incubated in a 37 °C shaker for 30 min. Cells were isolated by physical dissociation, filtered through a 70 µm cell strainer, and neutralized with cold DMEM. Samples were centrifuged at 350*g* for 5 min and resuspended in 500 µL cold FACS buffer (above). Cells were centrifuged at 350*g* for 5 min, resuspended in 1 mL cold FACS buffer with DAPI, pipetted up and down 5x gently with p1000, and strained through a 35 µm filter for FACS. Samples and cells were kept on ice throughout unless otherwise indicated.

### Liver and lung harvest and dissociation

To minimize blood contamination in the liver and lungs, 25 mL of cold PBS was perfused into the right ventricle of the heart (after blood draw from the heart). The entire liver (any macrometastases near the liver surface were completely excluded) and lungs were excised and processed identically to PTs, until immediately following the 30 min shaking digestion step. Here, samples were filtered through 100 µm cell strainers and then neutralized and centrifuged as with PTs, except 250*g* was used instead of 350*g* for centrifugation steps.

Liver samples were resuspended and further digested in 5 mL TrypLE for 5 min at 37 °C. Digestions were neutralized with cold DMEM + 10% FBS, centrifuged at 250*g* for 5 min, resuspended in 3 mL ACK, and incubated for 3 min at RT. Liver reactions were neutralized with cold PBS, centrifuged at 250*g* for 5 min, resuspended in 5 mL cold FACS buffer with DAPI, pipetted up and down 5 times gently with p1000, and strained through a 35 µm filter for FACS.

Lung samples were processed identically to liver samples except the order of ACK and TrypLE digestion steps was reversed (ACK before TrypLE). Additionally, lung samples were much smaller than liver samples and were thus only resuspended in 500 µL of cold FACS buffer with DAPI for FACS. Both liver and lung samples were kept on ice throughout unless otherwise indicated.

### Cancer FACS sorting and 10x Chromium loading

Cancer cells were isolated from dissociated tissues via FACS using a BD FACSAria II. After gating for singlets and live cells, GFP+ cells were sorted, thereby purifying PDAC cells from normal cells. For samples with a high yield of cells (PT, macrometastases, surgical-site), 30-35,000 cells were sorted on the purity setting. For each of the lung, liver, and blood samples, the entire sample was sorted on the yield setting to recover as many GFP+ cells as possible. The liver for M1 was stopped with 20% of the sample volume remaining due to excessively long sorting time. Cell numbers recovered for lung and liver were similar for each mouse (M1 liver: 22,000 (80% of total), M2 liver: 30,000, M1 lung: 1,000, M2 lung: 1,500).

After sorting, all samples were passed through a 30 µm filter and then centrifuged at 500*g* for 5 min and checked for visible pellets. Supernatant was removed to leave 20-30 µL of solution to not disturb the pellets. Remaining volume was measured and raised to 50 µL total by adding a 1:1 mixture of cold FACS buffer (without DNase) and nuclease-free water. 46.6 µL of these samples was loaded for 10x (Chromium Single Cell 3’ Reagent Kits v3), thereby superloading some lanes with up to 25-30,000 cells (macsGESTALT single cell barcode sequencing allows explicit detection of doublets, **Extended Data Fig. 10c, Methods: Clonal reconstruction and multiplet elimination**).

### scRNA-seq library preparation and sequencing

Single cell RNA-seq libraries were prepared as in the 10x Chromium Single Cell 3’ v3 user guide (Rev A) until Step 2.3. After cDNA amplification, the 100 µL cDNA PCR was split 50:50 for separate barcode and transcriptome library preparation. Transcriptome library construction continued as in the 10x user guide instructions. Indexed and pooled single cell transcriptome libraries for each mouse were sequenced separately on the NovaSeq 6000 System with S2 100-cycle kits.

### Single cell barcode library preparation and sequencing

For all single cell barcode PCRs (as for bulk DNA barcode PCRs), SYBR-containing master mix from the KAPA Real-Time Library Amplification Kit was used, and PCRs were stopped in mid-exponential phase. All primers were used at 10 µM. Primer sequence, purpose, and annealing temperature for all library preparation PCRs are included in **Supplementary Table 3**.

The barcode split of the cDNA amplification reaction (from 10x Single Cell 3’ v3 Step 2.2) was purified via 1.2x SPRI Select (Beckman Coulter #B23317). cDNA products were eluted in 40 µL of EB. Concentrations were measured by Qubit, and 2 ng/µL dilutions in EB were created for each sample. Barcode amplification and adaptor and sample index addition were performed in 2 sequential PCRs.

Barcodes were selectively amplified by PCR1. Here, 50 ng of each purified, diluted cDNA amplification sample was used to template a 100 µL PCR. After mixing, the reaction was split into 4 smaller reactions of 25 µL each for cycling. PCR cycling conditions were 1) 95 °C for 3 min, 2) 14-15 cycles of 98 °C for 20 s, 65 °C for 15 s, 72 °C for 15 s. Sample reaction splits were re-pooled after cycling, and products were purified with 0.9x SPRI Select and eluted in 60 µL EB.

Sample indices were added in PCR2. Here, 5-10 µL of the eluted products of PCR1 (1:12 or 1:6 overall dilution) were used to template a 100 µL PCR, which was again mixed and split into four smaller reactions of 25 µL each. PCR cycling conditions were 1) 95 °C for 3 min, 2) 6 cycles of 98 °C for 20 s, 65 °C for 15 s, 72 °C for 15 s. Sample reaction splits were re-pooled after cycling. Dual-sided size selection of complete barcode amplicons was performed using SPRI Select at an exclusion ratio of 0.5x and a selection ratio of 0.7x. Amplicons were eluted in 32 µL EB.

Barcode library size and concentration were checked via TapeStation HSD5000 and Qubit, respectively. Libraries were sequenced using Illumina MiSeq 600-cycle v3 Reagent Kits with the following run parameters: Read 1 - 28 cycles, i7 index - 8 cycles, Read 2 - 500 cycles. M1 was sequenced with 3 kits. Since barcode recovery only increased 5-10% with two additional kits for M1, M2 barcode library was sequenced with a single kit. Limiting dilution experiment libraries were also sequenced with a single kit.

### Single cell transcriptome data processing

Single cell transcriptome sequencing data was aligned and processed using 10x Cell Ranger v3. Filtered matrices from Cell Ranger output were further processed using Seurat 3.0 (https://satijalab.org/seurat/). All samples across both mice were merged into a single Seurat object. Low quality cells with ≤1,000 genes or ≥0.15 mitochondrial gene fraction (mito fraction) were filtered out. Cell cycle score and phase were determined for each cell using the CellCycleScoring function (https://satijalab.org/seurat/v3.1/cell_cycle_vignette.html).

Variable feature selection, scaling, and normalization were performed using SCTransform, while regressing cycle scores and mito fraction. Dimensionality reduction by PCA was performed using the first 15 principal components (PCs). Cells were plotted in UMAP space and a clearly-separated, large cancer cell cluster was observed, distinct from smaller clusters of contaminating normal cells, mostly derived from samples sorted on the FACS yield setting. Contaminating normal cells were filtered out. 10x cell barcodes, here referred to as cellIDs, for the cancer cells were then exported and used for initial macsGESTALT barcode data filtering.

### Single cell barcode data processing

Single cell barcode sequencing data was aligned, collapsed by UMI, and processed, as previously reported^4^ via a pipeline available on Github (https://github.com/mckennalab/SingleCellLineage/). For each sample, stats files, containing aligned and collapsed edited barcode sequence data, were extracted from pipeline output and used for clonal and subclonal analysis in R. Sample stats file for different harvest sites from a mouse were merged. However, each mouse and limiting dilution experiment was processed separately.

To ensure high-quality barcode data was used for reconstruction, five initial filtering steps were applied: First, cellIDs not present in the initial transcriptome cellID list (or v3 10x whitelist for limiting dilution experiments without transcriptional data) were filtered. Second, transcripts (UMIs) with incomplete static barcode (staticID) sequences were filtered. Third, staticIDs with less than two UMIs per cell were removed. Fourth, staticIDs with less than two UMIs per cell on average were filtered. Fifth, staticIDs found in less than 5 cells were filtered. Specific thresholds were determined by examining elbow plots of the relevant parameters.

### Clonal reconstruction and multiplet elimination

Next, potential clonal groupings of cells based on staticID content (absence or presence) were identified by complete-linkage hierarchical clustering. The staticID content of resulting clusters was examined, and clusters were found to be often improperly fractured due to cells with undetected staticIDs. To identify real clones defined by sets of staticIDs, clustering results were pruned by excluding clusters of less than five cells and staticIDs found in less than 20% of cells for a particular cluster. For clusters of less than 20 cells, staticIDs found in less than 35% of cells were further excluded. Then, clusters that were either duplicates or subsets of other clusters in terms of their defining staticIDs were collapsed. Finally, remaining staticID cluster sets were manually inspected for improperly fractured clusters, and any remaining improper cluster splits were merged or collapsed (usually this was either not necessary or was only needed for a few clusters).

After cluster cleanup, staticID sets were extracted and used to assign cells. Cells were matched to clusters based on their staticIDs. This process also served to explicitly identify interclonal multiplets, i.e. if a cell matched two or more clusters, this cell was removed as a multiplet. This method performed well, as only a small fraction of cells, ranging from 0 to 0.54% across experiments, went unmatched. Unmatched cells likely belonged to very small clones, only found in *in vivo* experiments. Furthermore, the percentage between mice was strikingly consistent (M1: 0.54% and M2: 0.51%), highlighting the reproducibility of the cancer model system and reconstruction approach. Only matched singlets were retained for downstream analysis.

With this orthotopic model, it is possible that some of the cells injected can leak out of the pancreas during and after injection and directly colonize the peritoneal cavity (although we sought to minimize this as previously described^17^). To eliminate any such cells from further analysis, we filtered clones that were detected in disseminated sites but not in the PT. This resulted in the removal of a small number of cells (M1: 1.49% and M2: 0%) from a few clones only found in peritoneal macrometases and in the surgical site lesion of M1.

In a true singlet, without genomic duplication of a barcode, each cellID-staticID pair should have a single mutagenized allele. To detect potential intraclonal multiplets or duplicated barcodes, we calculated the number of unique mutagenized evolving barcodes for a cellID-staticID pair, and mutagenized barcodes with less than 25% of the UMIs for that cellID-staticID pair were removed as technical noise.

PDAC is known to undergo large-scale copy-number changes via chromosomal instability. We observed this in our CNV analysis using InferCNV (**Extended Data Fig. 7**). While most staticIDs had a median of one mutated allele per cell, some had a median of two and a notably higher average. We speculated that these might be barcodes that resided in genomic areas that underwent copy number gain at some point after barcode integration. StaticID that had an average of 1.3 or greater mutated alleles per cell were considered to be potentially duplicated or triplicated.

Per 10x Chromium 3’ Single Cell v3 documentation (page 16), our overall expected multiplet rate for *in vivo* experiments with superloading was approximately 12% to 15%. Having explicitly detected and filtered interclonal multiplets, we next removed potential intraclonal multiplets. We filtered all cells with an average number of unique mutated alleles per staticID greater than 1.25, except for cells containing a potentially duplicated staticID; for these cells, the threshold was less stringent, at greater than 3. This resulted in appropriate overall multiplet rates of 12% for M1 and 15.7% for M2. Only true singlets were retained for further analysis.

After these filtering steps, clones that were detected in disseminated sites but not in the PT were again removed if present, and clones were then numbered by their size in the primary tumor, largest to smallest. These rankings are used to refer to clones throughout the paper with the mouse number appended, i.e. M1.1 or M2.14. These finalized clones were used for calculating clone size and clone fraction for each harvest site. These final filtered, clone-assigned singlets were used for further single cell transcriptional analysis.

Clonal aggression scores were estimated by giving points for size and fraction. For each non-PT harvest site where a clone was present 0.5 points were awarded. If the clone’s fraction was higher at a disseminated site than at the PT than it was rewarded an additional 1 point for that site. If a clone made up 5% or more of a disseminated site it received an additional 0.5 points for that site and a further 0.5 points if it was 10% or more.

For limiting dilution validation experiments, cells were visualized by their static barcode expression using tSNE in Seurat. A static barcode (rows) by cells (columns) expression matrix was generated. Just as in a regular transcriptome scRNAseq analysis, this matrix was used to generate a SeuratObject, which was then processed as a transcriptional dataset, where static barcodes were treated as features. The first 50 dimensions were used for tSNE plotting.

### Single cell transcriptional analysis

Transcriptional analysis continued using only singlets with quality barcode information (from above section). Seurat objects were converted into cell_data_set objects, and Monocle 3 (https://cole-trapnell-lab.github.io/monocle3) was used for all further transcriptional analysis. Preprocess_cds was run with top 20 dimensions (PCA) and align_cds was run with batch correction for harvest site and regression for cycle scores and mito fraction. Cells were plotted in UMAP space and two clusters of low quality or contaminating cells were removed. The first was a cluster of cells distinguished by high ribosomal fraction that was derived from cells of many clones and harvest sites. These cells were likely technical artifacts observed from droplet library preparation. The second was a cluster of cells with high hepatic gene expression. These cells derived from primarily the liver harvest sites and were most likely contaminating tumor-liver multiplets that had escaped initial filtrations steps.

Following these filtrations, preprocess_cds and align_cds were run again as before but with the top 25 dimensions, as determined by examining an elbow plot using plot_pc_variance_explained. Cells were plotted in UMAP space and clusters found using cluster_cells. Further transcriptional analyses and visualizations on all mouse cancer cells together were performed using Monocle 3 functions and custom R scripts as needed. For analyses on individual mice, cells were extracted and reprocessed as above but with the top 20 dimensions by PCA.

### Copy-number variation (CNV) analysis

InferCNV was used for single cell CNV analysis (https://github.com/broadinstitute/inferCNV/wiki). Default settings were used. Cutoff = 0.1 was used, which is recommended by InferCNV for 10x data. Clones were treated as cell groups, with cluster_by_groups = T. Clones with >200 cells were downsampled to 200. For clones ≤200 cells, all cells were included.

### PseudoEMT analysis

PseudoEMT or pseudotime analysis was performed by finding a trajectory in UMAP space using learn_graph with default settings. The root (most epithelial region) was placed where epithelial gene expression peaked. This additionally led to the most mesenchymal region existing at the end of the trajectory, thus resulting in a pseudoEMT spectrum. To find genes whose expression varied significantly along pseudoEMT, graph_test was used with the ‘principal_graph’ parameter selected. The top 3000 genes were retained, all of which had q ∼ 0 and Moran’s I > 0.1 (**Supplementary Table 1**). For the top 3000 genes, kinetic expression curves were clustered into groups by ward.D2 clustering using the R Pheatmap package, and the resulting tree was cut into six groups, which were named in order from epithelial to hybrid to mesenchymal patterns of expression.

To find enriched transcription factor motifs within the six gene clusters, findMotifs.pl from HOMER was used with the provided mouse promoter set. All default parameters were used, except for promoter region (−500, 50 bp from TSS) and background promoter frequency (derived from all top 3000 pseudoEMT genes). Known motifs passing an enrichment cutoff of p < 0.05 were extracted. The target genes of each motif were obtained using HOMER’s annotatePeaks.pl. Also for each pseudoEMT gene group, molecular signature database (mSigDB) gene set enrichment was determined using the hypergeometric test within HOMER.

### Subclonal and phylogenetic reconstruction

Using filtered barcode data (**Methods: Clonal reconstruction and multiplet elimination**), duplicated barcodes were removed entirely (this also removed any cells whose only recovered barcodes were duplicated). Cells with greater than one unique mutated allele per staticID were then filtered. For each cell in a clone, a barcode-of-barcodes was generated by concatenating all evolving barcode alleles, ordered by staticID. If a cell was missing a staticID, ‘UNKNOWN_UNKNOWN_UNKNOWN_UNKNOWN_UNKNOWN’ was concatenated for that staticID to note the missing information for all five target sites. Thereby, for an example clone defined by four staticIDs, every cell had four evolving barcodes concatenated in order and 20 target sites overall, including any missing information.

Within each clone, cells with identical barcode-of-barcodes were then grouped into subclones of indistinguishably closely related cells. To limit computational time required for downstream phylogenetic reconstruction of subclonal relationships, we pruned subclones of only a single cell from the largest clones, i.e. clones with ≥50 cells. This greatly increased computational efficiency while still retaining meaningful subclones.

Separate files were constructed for each clone, containing subclones with associated barcode-of-barcodes alleles. Phylogenetic reconstruction of subclonal relationships was performed for each clone barcode-of-barcodes file separately via TreeUtils (https://github.com/mckennalab/SingleCellLineage/). TreeUtils performed reconstruction using Camin-Sokal maximum parsimony via the PHYLIP Mix software package^55^ as previously described in depth^4^.

Further analysis then resumed in R. Clone Newick files were extracted from TreeUtils output and converted to an edgelist dataframe format. Clone edgelists were combined into a single large edgelist with a common root node (for each mouse separately). A small fraction of clones that were entirely defined by staticIDs that had been genomically duplicated, and were thus left out of phylogenetic analysis, were added back as a single node emerging directly from the root. At this point, cellIDs were added as terminal nodes emerging from subclone nodes (or directly to clone nodes for clones that were left out of phylogenetic analysis due to barcode copy gain). Cell nodes were then annotated with harvest site, transcriptional, and other information as needed. For circle pack or tree visualization, edgelist datafames were converted to igraph graph objects (https://igraph.org/r/) and plotted using ggraph (https://github.com/thomasp85/ggraph).

### Subclonal dissemination calculation

Shannon’s Equitability (E_H_) was used as a statistical measure of dissemination across harvest sites. To calculate E_H_, Shannon Diversity (H) was first calculated as follows:

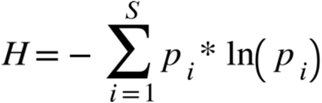

S is the number of distinct harvest sites analyzed (six for M1, four for M2). p is the sampling normalized proportion at which a subclone is recovered from a harvest site, i.e. if a subclone is only found in the PT, p_PT_ = 1, while p = 0 for all other sites. A subclone’s H is then used to calculate its E_H_ as follows:

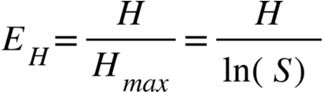

E_H_ therefore normalizes H by the number of harvest sites analyzed to exist between 0 and 1, with 1 being completely even dissemination and 0 being no dissemination. For example, a subclone found at only one harvest site is not metastatically aggressive and has an E_H_ = 0.

### PseudoEMT across ancestral relationships

Comparison of pseudoEMT for root clades, subclones, and cells was performed in R. To determine root clade pseudoEMT values, we recursively calculated the weighted mean pseudoEMT value of ancestral nodes moving backwards along phylogenetic trees. Root clades were the nodes immediately preceding the common root of M1.1. These clades are depicted by the outermost circles in the circle packing visualizations of M1.1 (**Fig. 4a**,**b**). The density of root nodes, subclones, and cells along the pseudoEMT axis was then plotted as a ridge plot for comparison.

### PseudoEMT gene cluster TCGA survival analysis

PseudoEMT genes (n = 3000) were mapped to their human homologs using getLDS() from the biomaRt package. All homologous genes were included. Preprocessed transcriptomic data (FPKM abundance after upper quantile normalization; FPKMuq) for pancreatic adenocarcinoma patients (n = 173) from the TCGA Pancreatic Adenocarcinoma project (TCGA-PAAD; https://www.cancer.gov/tcga) were obtained using the R package TCGAbiolinks. Using the singscore^56^ package, patients’ enrichment scores were determined for each pseudoEMT gene cluster (E, H1, H2, H3, H4, M). Survival (from the time of pathological diagnosis) was obtained from TCGA-PAAD clinical data. Univariate and multivariate Cox regression analysis was performed in the R environment (survival, survminer) to determine the hazard associated with each pseudoEMT gene signature. Wald test, LLR and Score test were all significant, indicating the regression model was significant (p = 0.02).

### Pseudobulk and metagene analyses

The aggregate_gene_expression function from Monocle 3 was used to perform pseudobulk and metagene analyses. For testing whether clones retained their transcriptional identity, pseudobulk samples consisting of clone and harvest site combinations were generated, and only pseudobulk samples with >20 cells were used for further analysis. The entire transcriptome for each pseudobulk sample was aggregated and used to hierarchically cluster samples via the Pheatmap package, with the ward.D2 clustering option.

### Data availability

Single cell lineage tracing data and transcriptional data will be made available online via a data sharing platform, such as Dryad, figshare, or Gene Expression Omnibus. Plasmids needed to implement macsGESTALT will be deposited in Addgene.

### Code availability

Code used for clonal and subclonal reconstruction and related analyses will be made available via GitHub.

## Acknowledgements

We thank J.I. Murray for advice on lineage and transcriptional analyses, J. Li for donation of PDAC cell line and advice on its use, J.A. Gagnon for advice on barcode editing and lineage tracing, and M.A. Blanco for advice on TCGA survival analysis. We thank K. Tan and A. Raj as well as all members of the Lengner laboratory for helpful discussions. We also thank the University of Pennsylvania Next-Generation Sequencing Core, in particular J. Schug and J. Kutch, for advice on barcode sequencing. We thank D.P. Beiting for computational resources. This research was supported by the Ruth L. Kirschstein National Research Service Award F30-DK120135, the Blavatnik Family Fellowship in Biomedical Research, and T32-HD083185 (to K.P.S.), National Cancer Institute R01-CA168654 and the Shipley Foundation Program for Innovation in Stem Cell Science (to C.J.L.), and the Howard Hughes Medical Institute and Allen Discovery Center for Cell Lineage Tracing (to J.S.).

## Author contributions

K.P.S. initiated, designed, and coordinated the study with the guidance of A.M., J.S., and C.J.L.; K.P.S., B.M., and A.M. constructed vectors; K.P.S. generated cell lines and performed in vitro experiments; R.J.N. performed orthotopic injections and advised on the PDAC model with B.Z.S.; K.P.S. and M.L.C. harvested and isolated tumor cells; K.P.S. performed bulk and single cell library preparation and sequencing, wrote clonal reconstruction and subclonal analysis scripts, and performed all single cell lineage and transcriptional analyses; C.N.B. performed motif enrichment, copy-number variation, and survival analyses; K.P.S. wrote the manuscript and generated all figures and data visualizations; K.P.S., C.N.B., M.L.C., A.M., J.S., and C.J.L. reviewed and edited the manuscript.

---

**Extended Figure 1:**
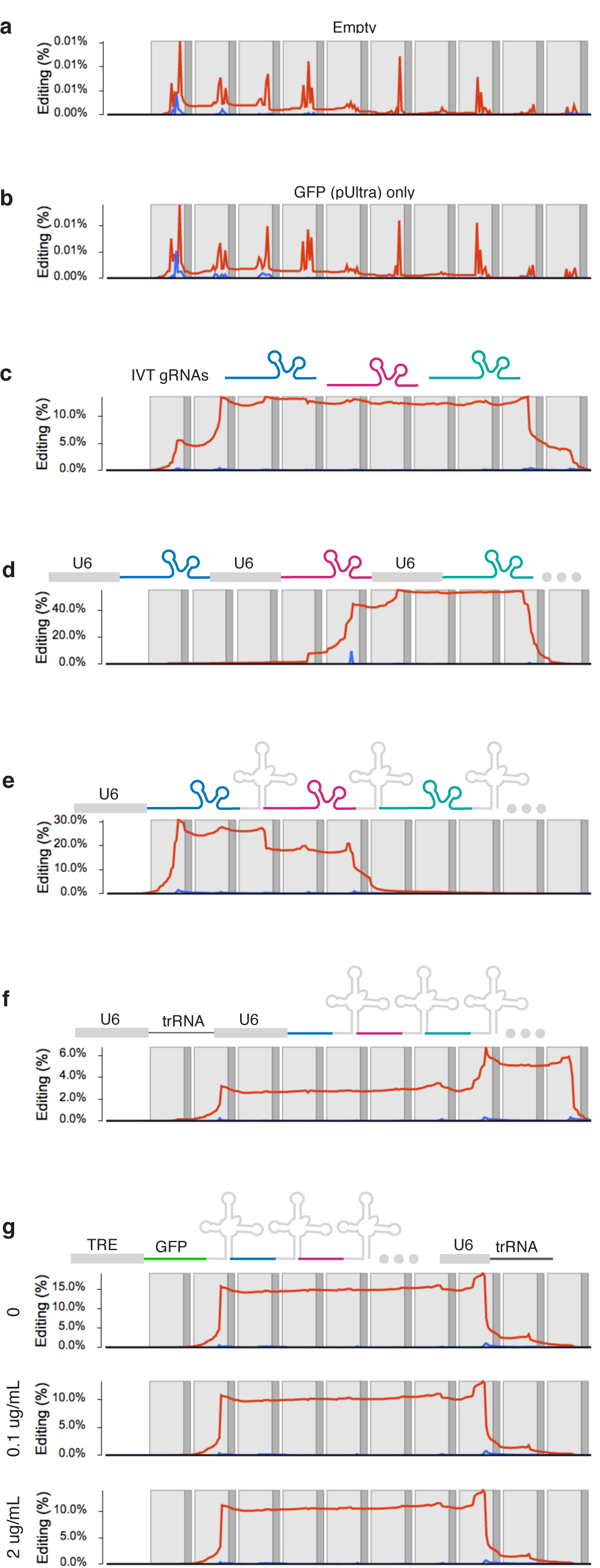
gRNA array editing screen. Barcoded 293T cells were transfected with constitutive Cas9 vector (px330) and co-transfected with a variety of controls or gRNA expression formats. Barcode genomic DNA was collected and bulk sequenced one week post-transfection, and for each condition, the percent at which each barcode base was deleted (red) or adjacent to an insertion (blue) is indicated, along with target site spacers (light grey) and PAMs (dark grey). Controls included no transfection (only Cas9) negative control (**a**), GFP-only negative control (**b**), IVT gRNAs positive control (**c**), and each gRNA placed (targeting sites 5-9) under its own U6 promoter (**d**). **e**, gRNA-tRNA array (targeting sites 1-5) under a U6 promoter (selected for PDAC experiments due to both high editing rate and compact size). **f**, A split gRNA array with a crRNA-tRNA portion (targeting sites 2-10) and a tracrRNA portion under U6 promoters, where the crRNA portions can complex with the tracrRNA portions when expressed. **g**, The same array is in **f** but with the crRNA-tRNA array in 3’UTR of a dox-inducible GFP and cultured in three different doses of dox post-transfection for 5 d (this configuration was leaky with no change in editing rate with dox administration).

**Extended Figure 2:**
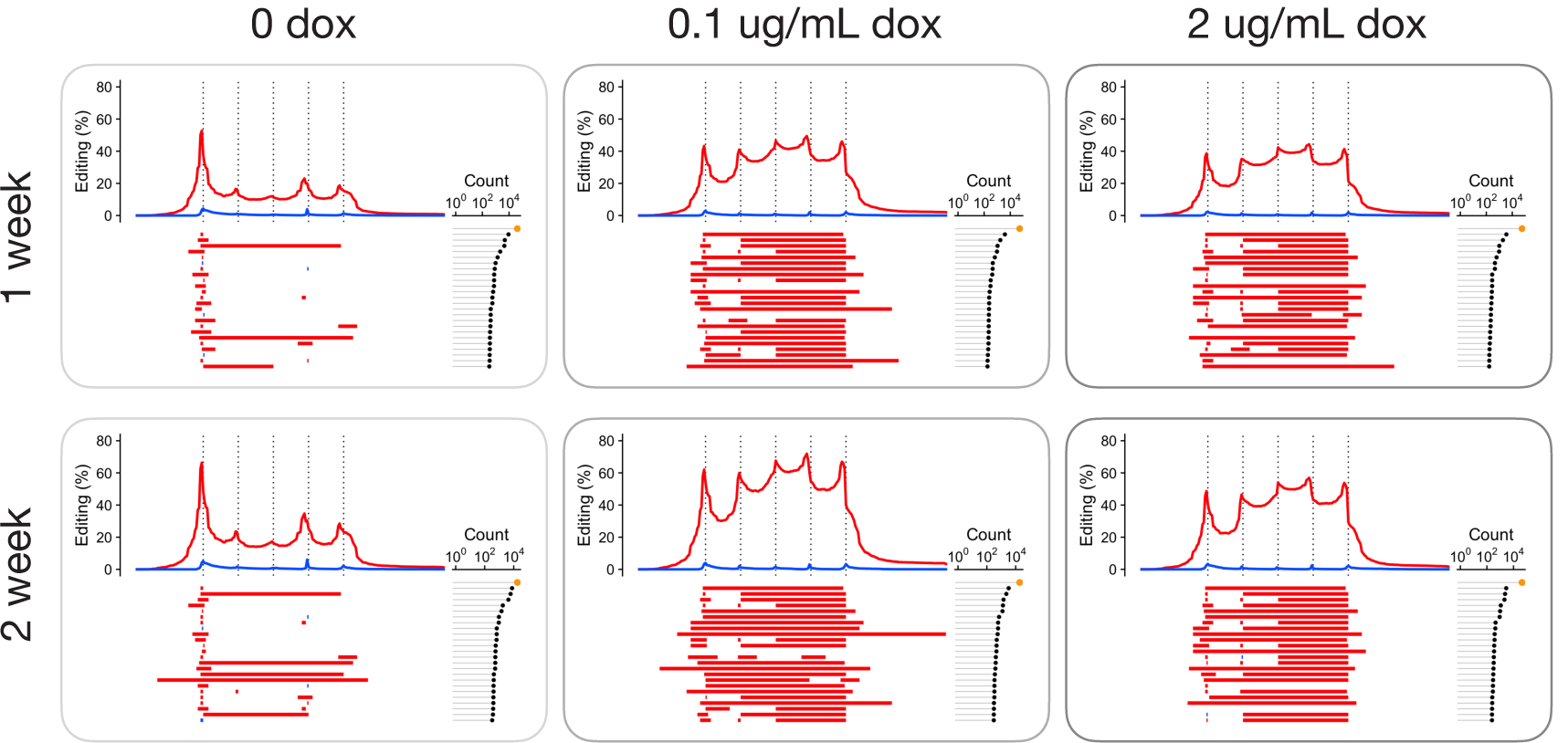
Dox-induced macsGESTALT PDAC cells edit evenly across sites and accumulate edits over time. macsGESTALT PDAC cells cultured in dox for one (top) or two (bottom) weeks, and barcodes were bulk DNA sequenced. The percent at which each barcode base was deleted (red) or adjacent to an insertion (blue) is indicated, along with expected cut sites (dotted lines, 3 bp upstream of PAMs). Beneath editing plots, the top 25 most commonly observed alleles are illustrated with the number of observations for each on the right. Leakiness was primarily localized to the first target site, while sites 2-5 remain largely unmutated until dox administration.

**Extended Figure 3:**
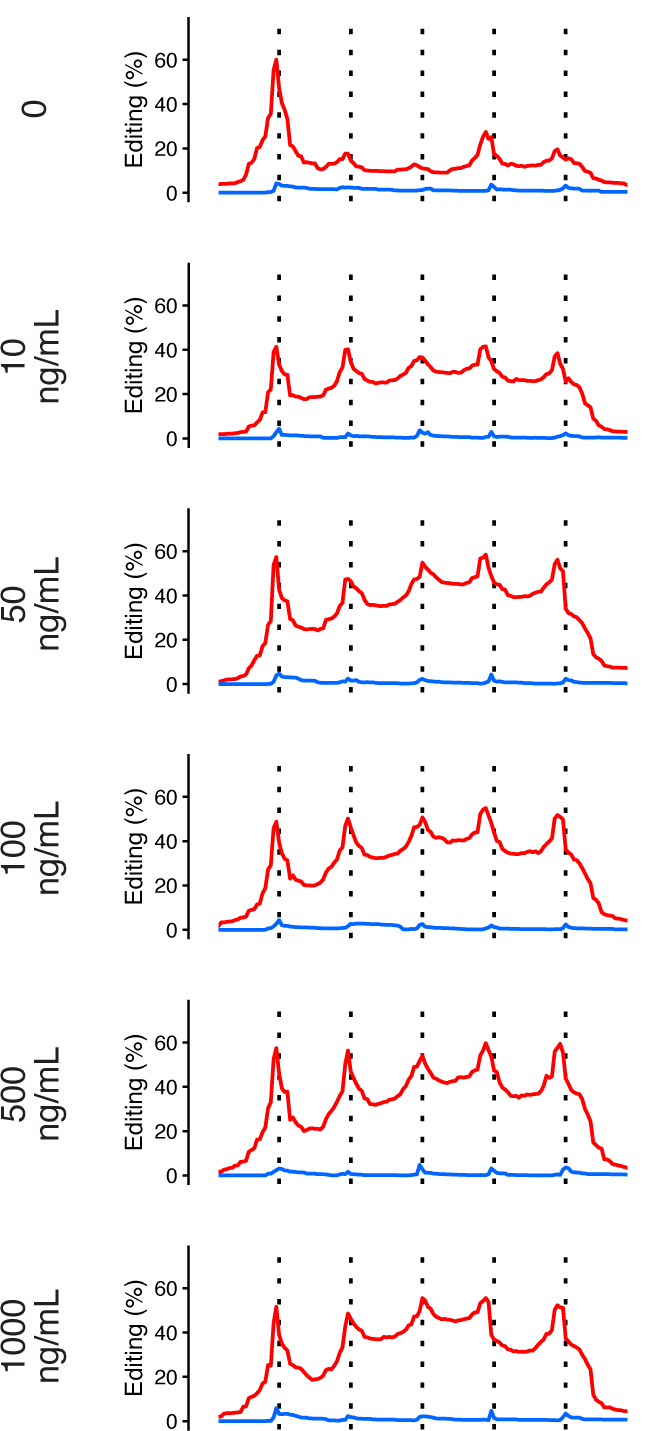
Dox-inducible editing initiates and peaks at low doses in macsGESTALT PDACs. Cells were cultured under six different dosages of dox for 2 weeks and barcodes were bulk DNA sequenced and editing rates plotted.

**Extended Figure 4:**
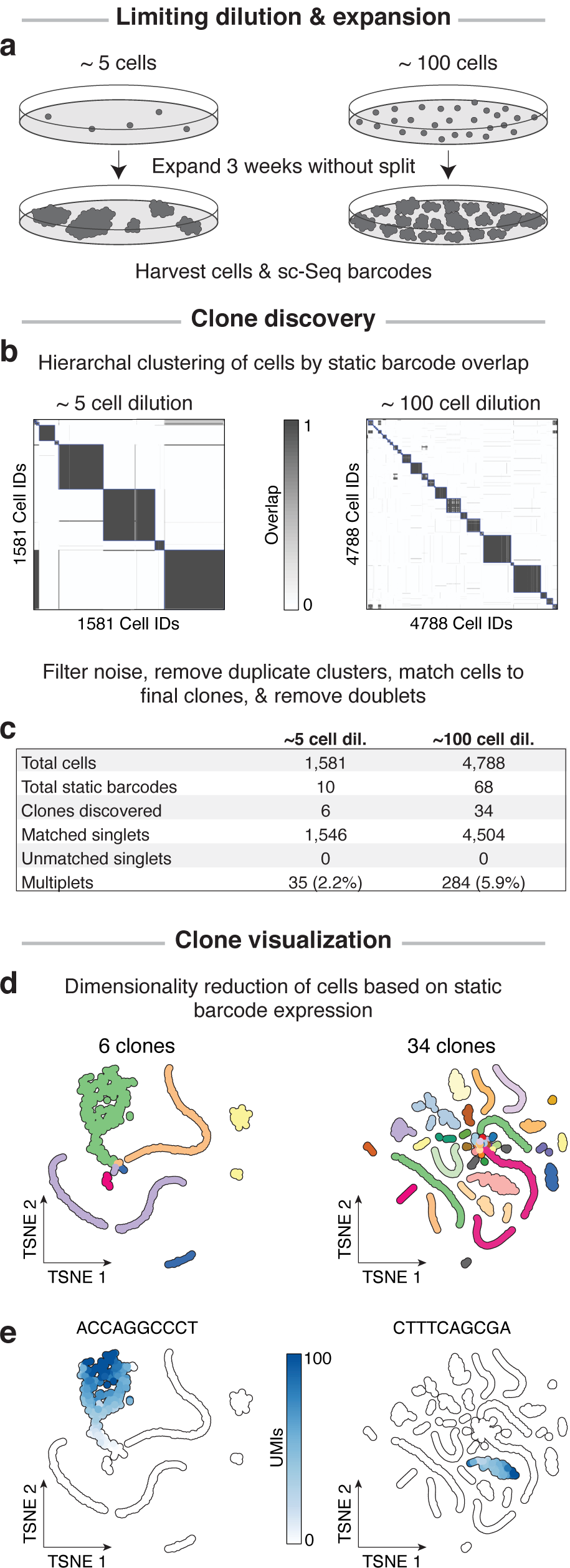
*In vitro* validation of clonal reconstruction and single cell readout of macsGESTALT PDAC cells. **a**, macsGESTALT PDAC cells were plated at two limiting dilutions, expanded without splitting (even if confluent), and barcodes were scRNA sequenced. **b**, Potential clones of expanded cells were identified based on static barcode overlap via hierarchical clustering. **c**, Approximately the expected number of clones were identified for the 5-cell dilution, while the 100-cell dilution retained a smaller fraction of clones likely due to extended culture time under confluence. All cells were successfully matched to a clone and multiplets were explicitly identified (**Methods: Clonal reconstruction and multiplet elimination**). **d**, Cells were plotted in tSNE space based on their static barcode expression. Cells clustered in tSNE space consistently with their clonal assignments. **e**, Two examples illustrating static barcode expression confined to specific clones.

**Extended Figure 5:**
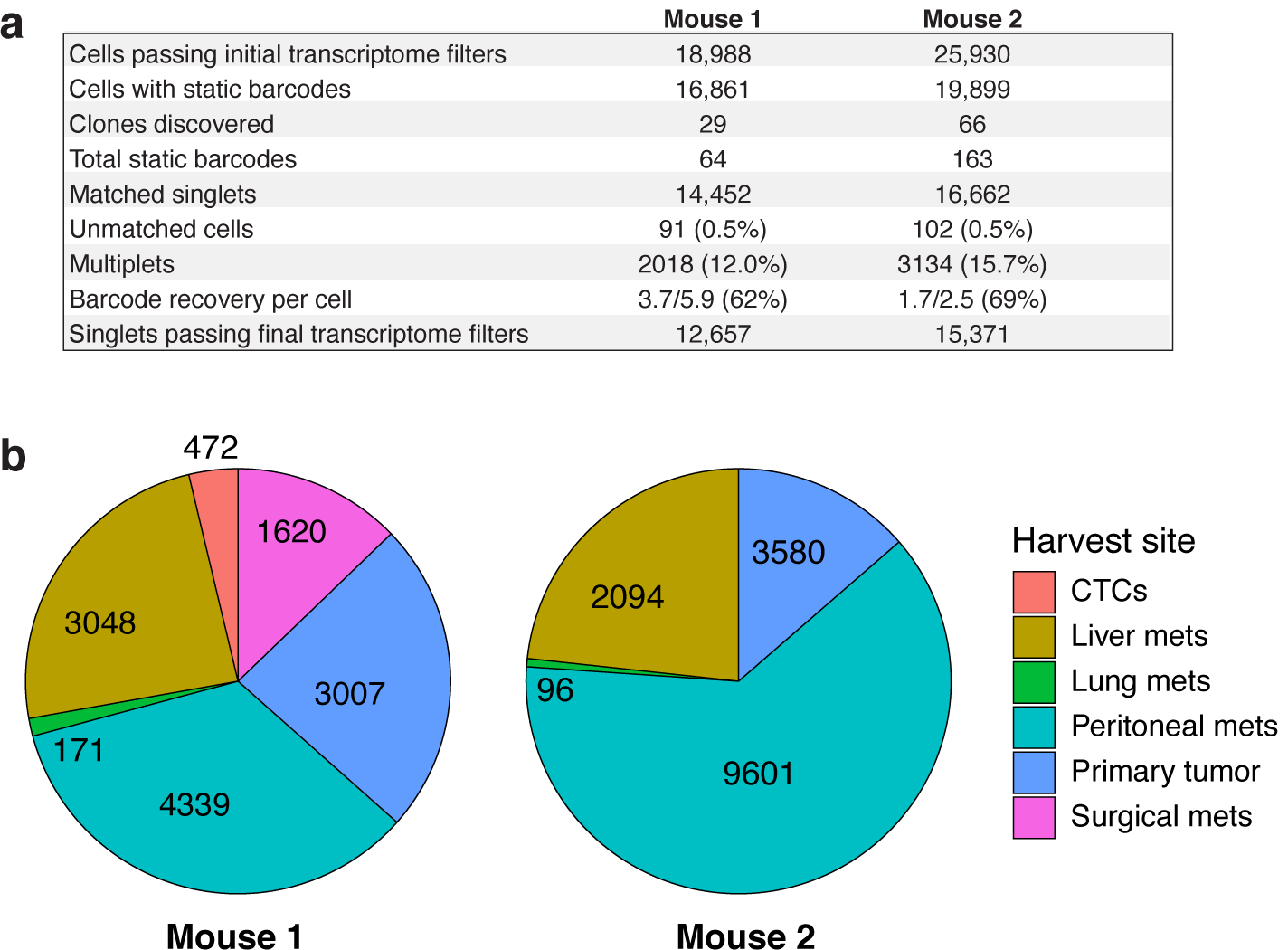
Summary of barcode, cell, and clone recovery in metastasis experiments. **a**, Table summary for each mouse. **b**, Number of single cancer cells obtained from each harvest site after filtering.

**Extended Figure 6:**
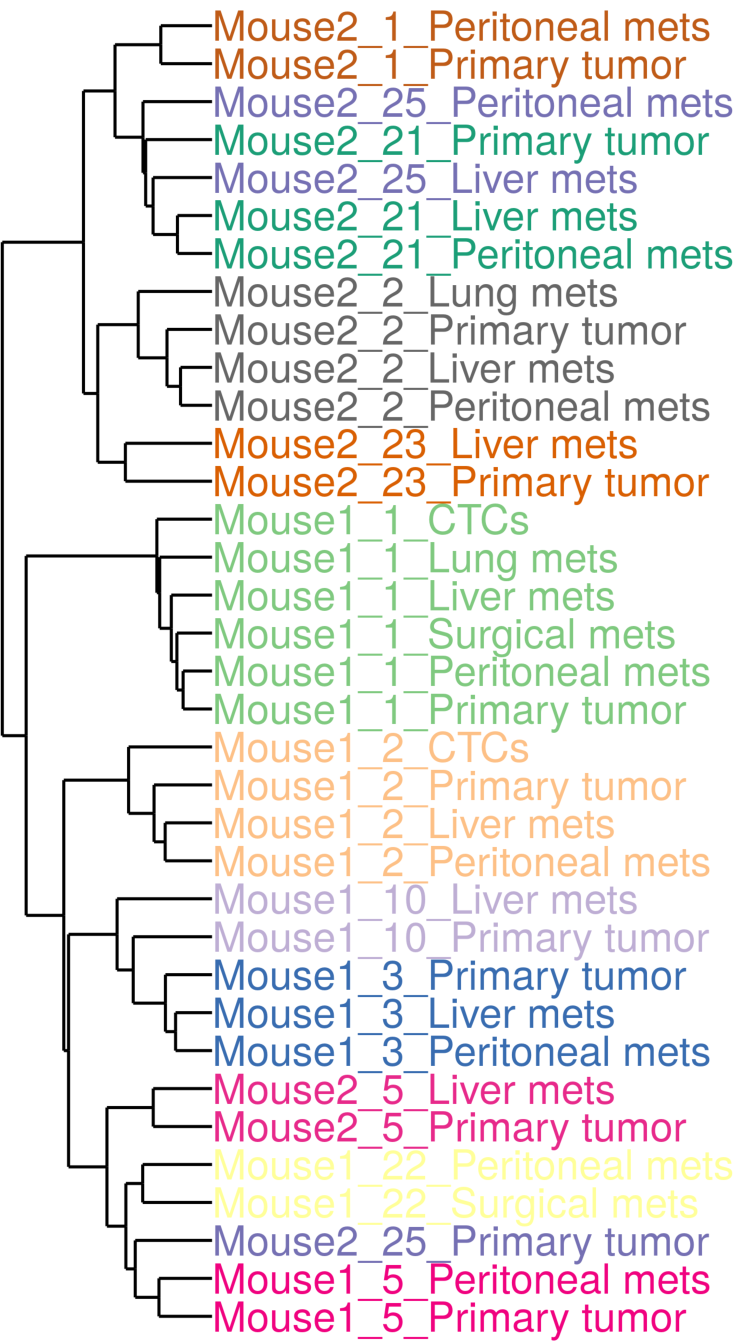
Clones retain transcriptional identity after metastasizing. Cells were analyzed as clone-site pseudobulk samples (i.e. cells from each clone and harvest site combination were aggregated and treated as a bulk sample, **Methods: Pseudobulk and metagene analyses**) and each sample was colored by clone. Only pseudobulk samples with >20 cells were used. Clone-site pseudobulk samples were hierarchically clustered based on whole transcriptome expression. Pseudobulk samples displayed preferential clustering by clone rather than harvest site.

**Extended Figure 7:**
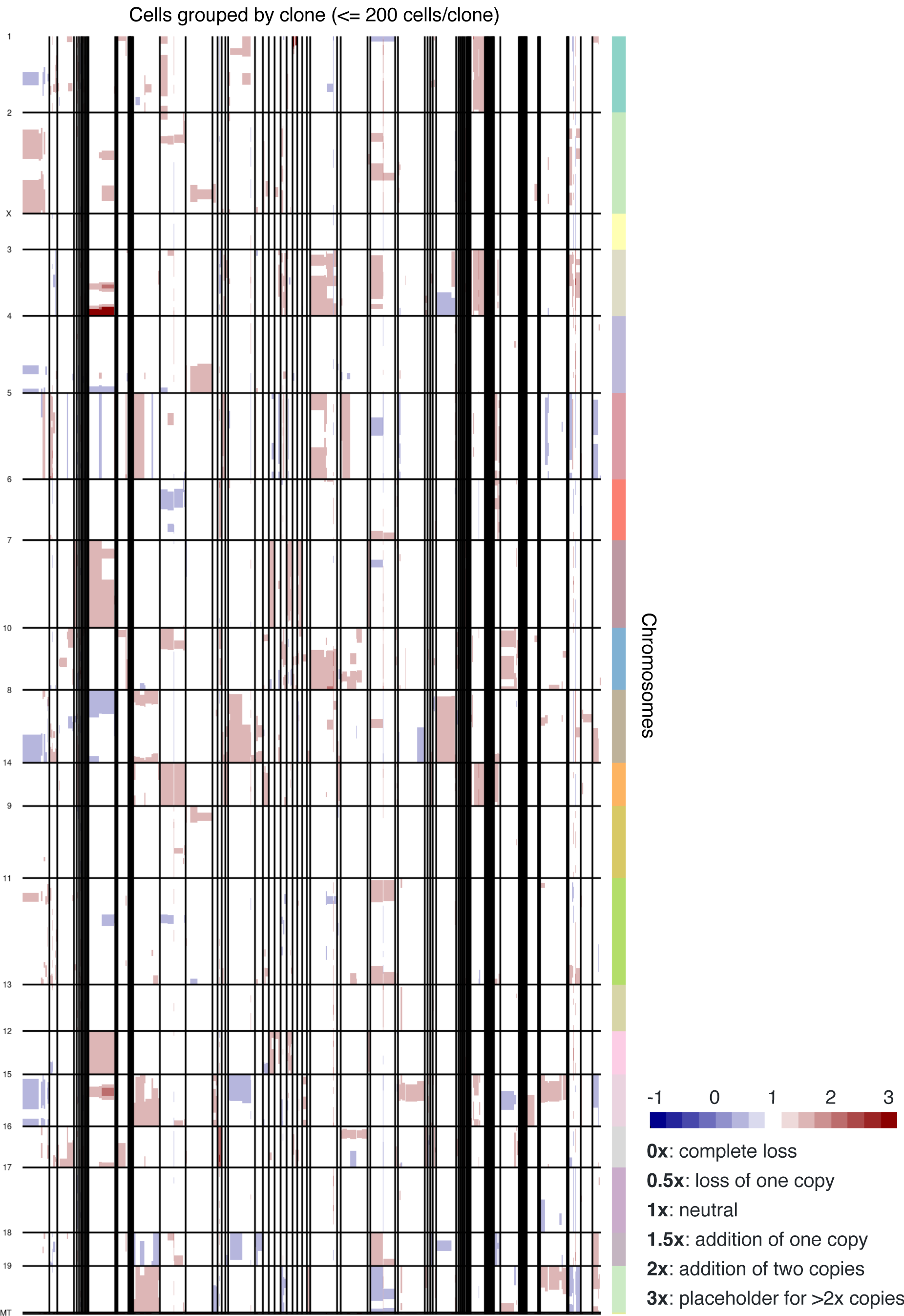
Genomic copy-number changes among clones. Copy number variation analysis was performed on all 95 clones. Clones with >200 cells were downsampled to 200 cells to perform CNV analysis with InferCNV. Vertical black lines divide clones (many small clones are not visible), and horizontal lines divide chromosomes. Large scale copy number changes are visible between and within clones.

**Extended Figure 8:**
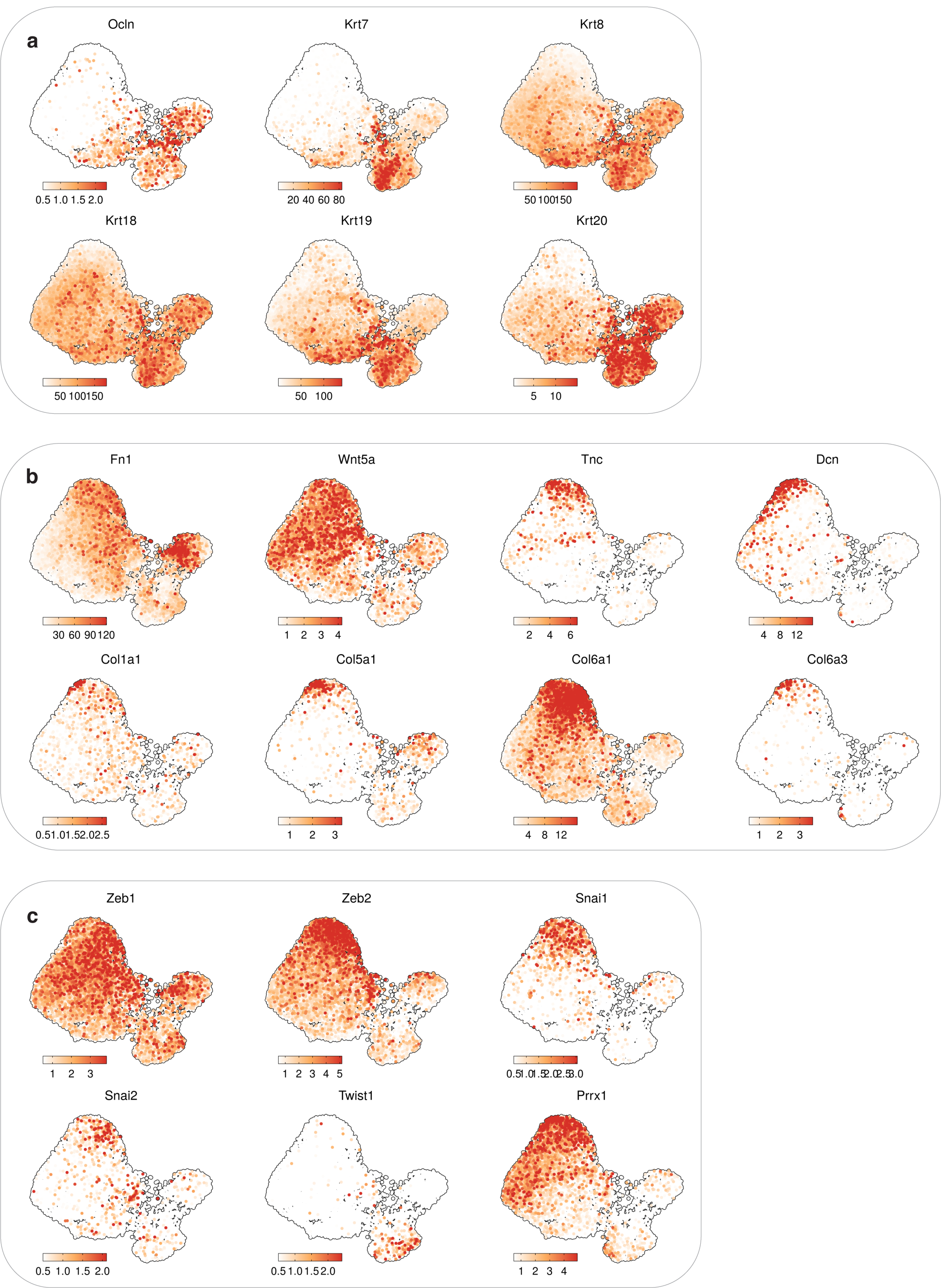
Expression of epithelial and mesenchymal genes. Epithelial markers (**a**), mesenchymal markers, including extracellular matrix genes (**b**), and canonical EMT-TFs (**c**) expressed in M1 cells.

**Extended Figure 9:**
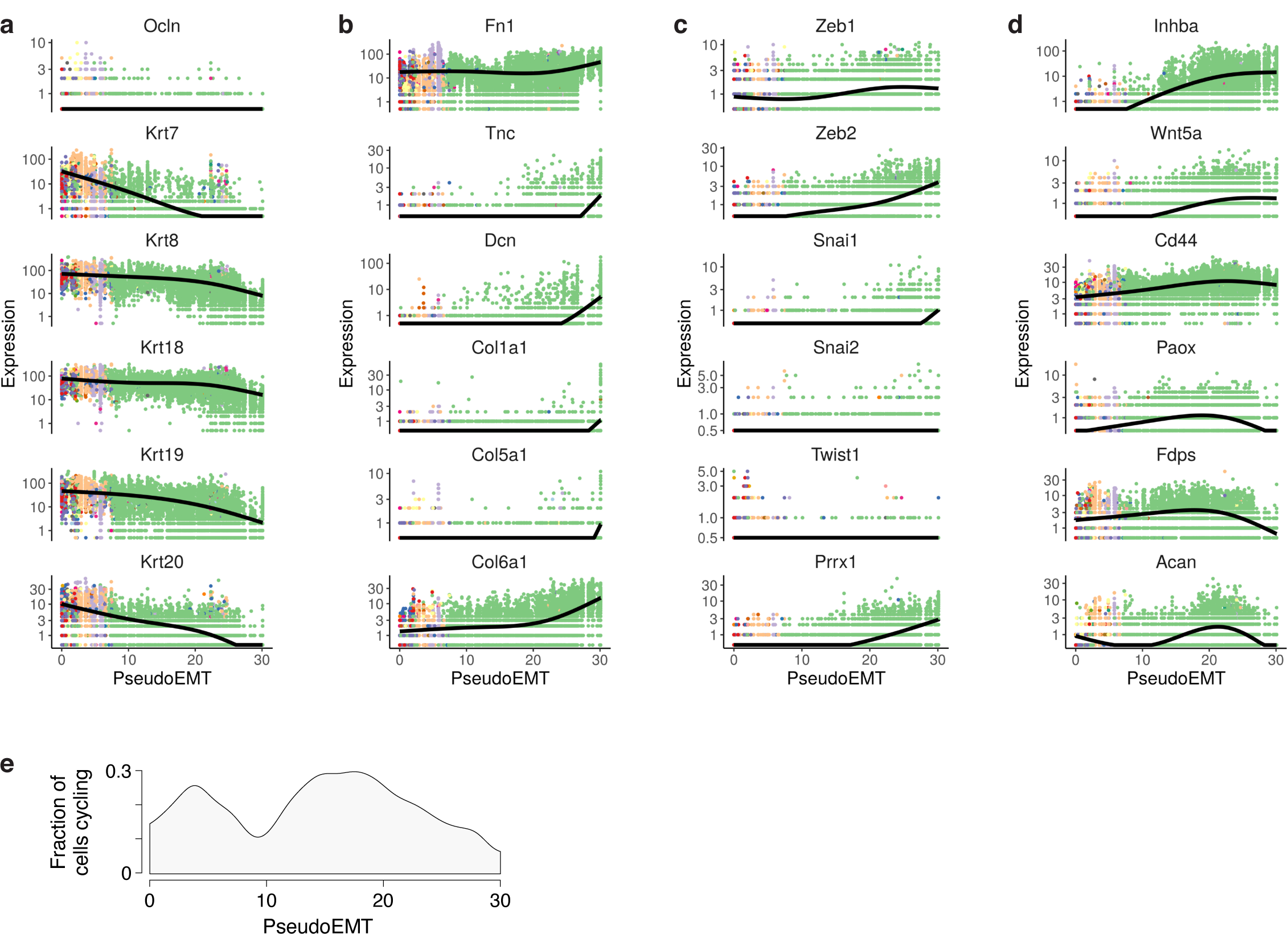
Fluctuation of genes and cell cycling across pseudoEMT. Epithelial markers (**a**), extracellular matrix mesenchymal genes (**b**), canonical EMT-TFs (**c**), and selected genes with unusual kinetics (**d**) across pseudoEMT. **e**, Fraction of cells cycling, i.e. in S/G2M cell cycle phase, across pseudoEMT.

**Extended Figure 10:**
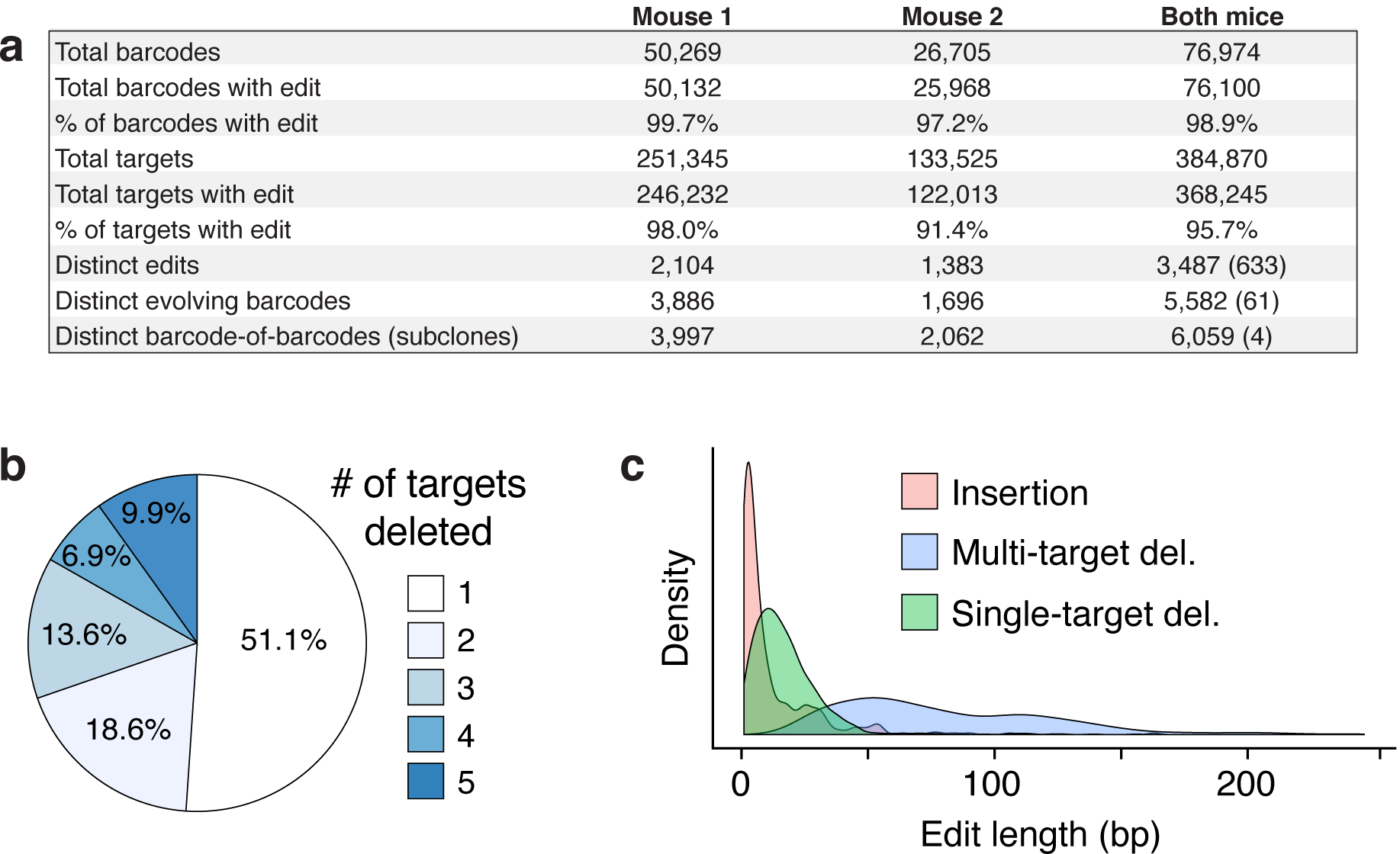
Editing summary of evolving barcodes in metastasis experiments. **a**, Summary table of the number of barcodes/target sites recovered, and the rate at which they were observed to carry a mutation. Additionally, the number of distinct edits, evolving barcodes, and barcode-of-barcodes are displayed. In the last three rows in the last column, the number of overlapping edits, evolving barcodes, and barcode-of-barcodes between the mice is indicated in parentheses. **b**, The proportion at which a deletion impacts 1, 2, 3, 4, or 5 target sites. **c**, Size distribution for insertions, single-target deletions, and multi-target deletions.

